# Barely depictive: Predicting imagery vividness relative to perception with EEGNet

**DOI:** 10.64898/2026.03.11.711041

**Authors:** Claire Vanbuckhave, Giorgio Ganis

**Affiliations:** Faculty of Health, School of Psychology, University of Plymouth, UK; Brain Research and Imaging Centre (BRIC), Faculty of Health, University of Plymouth, UK

**Keywords:** visual mental imagery, visual perception, vividness, convolutional neural networks, EEG

## Abstract

Previous studies suggest that visual mental imagery (VMI) acts as a weaker form of top-down visual perception (VP), with the two becoming more similar as VMI vividness increases. However, this relationship remains ill-defined, and it is unclear precisely how much weaker VMI is relative to VP. Here, we introduce an original probabilistic deep learning approach to quantify vividness at the neural level. Thirty-four participants either imagined or perceived stimuli presented at varying levels of vividness and provided trial-by-trial, picture-based vividness ratings. EEG activity recorded during VP was used to train a convolutional neural network (EEGNet) to predict perceived vividness from eight posterior electrodes located around early visual areas. A leave-one-subject-out cross-validation procedure showed that the model generalised across participants with above-chance accuracy during VP. On VP trials, predictions tracked vividness labels, with reliable interpolation to new vivid labels not included during training. Applied to VMI trials, mean expected VMI vividness remained substantially lower than expected vividness for seen stimuli but slightly higher than baseline, supporting a ‘barely’ rather than ‘quasi’ depictive imagery. For 91% of participants, mean expected VMI vividness was also lower than, yet scaled with, mean reported VMI vividness. This framework provides a principled way to quantify and compare VMI and VP on a shared neural-behavioural scale, with implications for studying individual differences and aphantasia.

## 1. Introduction

Whenever we look at something, our brains transform the visual information conveyed by the corresponding sensory input—such as its shape, colour, orientation, contrast, brightness, etc.—into complex patterns of neural activity. Evidence for this comes from the fact that many properties of this visual information, ranging from low-level features to higher-level category information, can be decoded from the neural activity patterns elicited by seeing a given stimulus (see Wilson et al., 2024, for a review). Part of the visual information initially encoded during visual perception (VP) can still be retrieved and experienced in the absence of the corresponding sensory input, an ability known as visual mental imagery (VMI; Pearson et al., 2015). Research has shown that the brain treats visual information originating from VMI (the mind’s eye) and VP (the real eyes) in a relatively similar way (Pearson & Kosslyn, 2015; Slotnick et al., 2005). For instance, training a classifier on VP and testing it on VMI has revealed shared neural representations of simple patterns in early visual areas (Huang et al., 2023a), as well as stimulus identity (Ragni et al., 2020) and category (Kilmarx et al., 2024). Although reduced patterns of activations are generally observed during VMI compared to VP, the neural overlap between the two tends to increase when people report experiencing more vivid or ‘perception-like’ imagery (Dijkstra et al., 2017).

Reported VMI experiences are often described on a continuum of vividness, from absent to as clear as real seeing (Zeman et al., 2020). Yet, many findings support the idea that VMI generally is more similar to ‘a weaker form of top-down VP’ at the neuro-behavioural level (Kosslyn et al., 2006; Pearson, 2019; Pearson et al., 2015; but see: Arcangeli and Bartolomeo, 2025; Cavedon-Taylor, 2021). The claim of a reversed cortical information flow between VMI and VP (‘top-down VP’) has been supported by direct comparisons between VP and VMI using high-density EEG (Dentico et al., 2014). However, the relative vividness of mental images compared to VP (‘weaker form of’) remains to be quantified at the neural level. In fact, beyond subjective reports, there is limited insight into vividness as a perceptual property of VMI itself.

Cross-task decoding analyses of neural data, that is, training an algorithm to decode perceptual content during VP and testing it on VMI, offer a promising tool to investigate this matter. However, to our knowledge, no study has yet attempted to cross-decode vividness between VP and VMI. The likely reason is practical: decoding and cross-decoding require labels. Category labels (e.g., ‘face’, ‘house’) can be externally controlled and have a verifiable ground truth in both tasks, but what about vividness labels?

The first step in obtaining vividness labels is to clearly define and operationalize vividness. To enable meaningful VMI–VP comparisons, this operationalization must be consistent across tasks (VMI and VP) and across the *loci* or levels at which vividness is investigated. Here, we distinguish vividness as (1) a property of the visual information conveyed by the stimulus itself (stimulus level), if any; (2) a property of that information as encoded in neural representations (neural level); or (3) a property of the resulting visual experience, if any (phenomenological level). Recent literature reviews suggest that phenomenological vividness can be operationalized along two main perceptual dimensions: clarity and intensity (Fazekas, 2024; Marks, 2023). At the stimulus level, this corresponds to variations in the physical properties of the input stimulus, such as its sharpness, intensity, or contrast. Accordingly, in perception tasks, neural-level vividness can be investigated by manipulating these low-level stimulus features and measuring corresponding brain signals.

In imagery tasks, however, obtaining reliable and objective vividness labels is inherently more challenging. Indeed, we cannot manipulate or control how vividly a given stimulus is being imagined: we can only measure it. Because mental images are, by definition, confined to one’s mind, current measures of imagery vividness must rely on introspection (e.g., via trial-by-trial ratings or questionnaires) or indirect markers (e.g., pupil size, perceptual priming). While there is some evidence to support the validity of such measurements (Arnold et al., 2024; Cui et al., 2007; Kay et al., 2022; Keogh & Pearson, 2018), they have recently yielded mixed results (Azañón et al., 2025; Bouyer et al., 2025; Gardner et al., 2026; Vanbuckhave et al., 2026) and correlations with neural activity remain generally weak to moderate at best (Bergmann et al., 2016; Cabbai et al., 2024; Huang et al., 2023b; Runge et al., 2017).

To overcome these limitations, the present article introduces a novel approach to assessing the relative vividness of VMI and VP on a common neural-behavioural scale. The validity of the approach rests on four main evidence-based assumptions. First, vividness can be operationalised as a combination of low-level visual features, such as clarity and intensity (Fazekas, 2024; Marks, 2023). Second, neural representations in early visual areas (EVAs) carry information about these low-level features during both VMI and VP (Naselaris et al., 2015; Wilson et al., 2024). Third, neural representations in EVAs are at least partially shared between VMI and VP (Dijkstra, 2024; Huang et al., 2023a; Pearson & Kosslyn, 2015; Slotnick et al., 2005). Fourth, these neural representations can be measured with electroencephalography (EEG) and their low-level features can be reliably decoded with machine learning or deep learning methods. This fourth assumption is implicit in most of the studies cited above.

Given these assumptions, the proposed approach exploits neural activity elicited during VP to generate probabilistic predictions about vividness during VMI. In other words, neural signals that successfully encode the vividness of perceived stimuli are used as a reference to estimate the expected vividness level when these same stimuli are imagined. In practical terms, this framework addresses the following question: given the neural evidence used to classify vividness during perception, what relative level of vividness would be predicted when the same stimuli are generated through imagery?

To address this question, we recorded EEG while participants imagined or perceived stimuli presented at different levels of vividness. The two tasks were designed to maximize similarity (timing, instructions, response format), and participants provided trial-by-trial, picture-based vividness ratings during both tasks. To extract a subject-independent neural signature of vividness, a convolutional neural network (CNN) was trained to decode operationalized vividness across participants from single-trial EEG activity in EVAs during the perception task. The trained model was then applied to imagery data, projecting unlabelled VMI trials onto the perceptual vividness axis learned from perception. Finally, the model’s predictions for VMI were compared with both its predictions during perception and participants’ reported imagery vividness.

## 2. Methods

### 2.1. Participants

Forty participants initially completed the experiment. Five were excluded due to technical issues during data collection (e.g., recording failures, data loss), and one participant was excluded for excessive errors during the perception phase (i.e., clear outlier; mean accuracy of 54.3% which was lower than the overall mean accuracy by more than 3 standard deviations), suggesting possible confusion or non-compliance to the task. The final sample therefore comprised 34 participants (age: *M* = 19.8, *SD* = 1.1, range 18-23), of whom 18 identified as female (age: *M* = 19.9, *SD* = 1.2), 15 as male (age: *M* = 19.7, *SD* = 1.0), and one did not report their gender.

### 2.2. Materials

The experiment was implemented in PsychoPy (v2024.2.2) with the visionscience package (v0.0.7) to present dynamic visual noise. Stimulus processing and adjustments were performed in MATLAB (R2024b) using the Image Processing Toolbox (v24.2). All analyses were carried out in Python (v3.11.13), with the sklearn (v1.7.1), MNE (v1.11.0), torch (v2.10.0) and tensorflow (v2.20.0) libraries.

All EEG data were recorded at 512 Hz using the BioSemi ActiveTwo system (ActiView software) with 64 electrodes arranged according to the 10-10 standard. Stimuli were displayed on a 50.8 × 35.9 cm ViewSonic VX2268WM monitor (120 Hz refresh rate) positioned 120 cm from participants. All sessions were conducted in a dimly lit room.

### 2.3. Stimuli

The visual stimuli consisted of two coloured images obtained from Google Images (a cat and a strawberry) and presented on a uniform dark grey background (RGB = [0.2, 0.2, 0.2]). Stimulus vividness was operationalized by parametrically degrading two low-level visual properties: spatial sharpness and luminance contrast, as proxies for clarity and intensity variations (see Introduction).

To manipulate sharpness, each image was filtered using a two-dimensional Gaussian smoothing kernel with standard deviations of σ = 2, 10, 20, and 40, applied identically to all three RGB channels. Increasing σ resulted in progressively stronger spatial blur. To manipulate luminance contrast, pixel values in each RGB channel were linearly scaled by multiplicative factors of 1.00, 0.75, 0.50, or 0.25 and blended with the grey background, thereby reducing contrast relative to the display background rather than uniformly darkening the image (see Data Availability section for the code). The highest vividness level (level 5) corresponded to minimal blur (σ = 2) and full intensity (1.00), whereas lower vividness levels (levels 2, 3 and 4) combined increased blur and reduced contrast. The lowest level (1) consisted of the dark grey background alone (no image; control/baseline condition). Figure 1 illustrates the response options presented at the end of each trial, with the five vividness levels for both stimulus categories.

**Figure 1.**
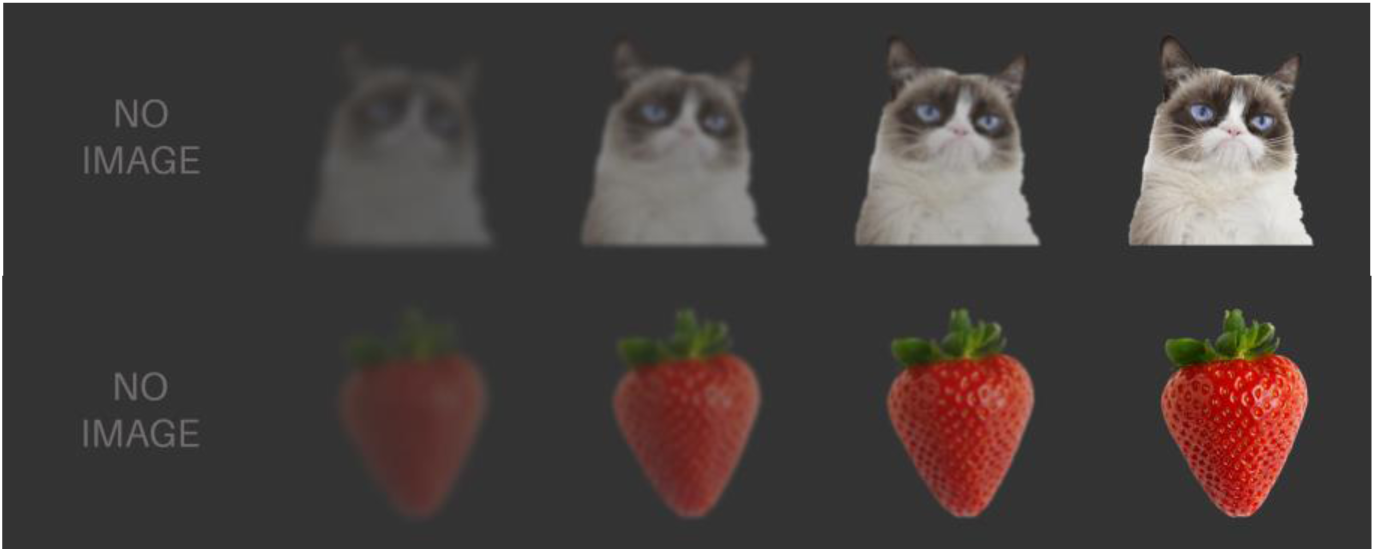
Response options for the two stimuli and their five vividness variations. Responses options corresponded to all possible vividness levels shown during the perception task, except for the ‘no image’, which corresponded to the dark gray screen alone (without text).

### 2.4. Procedure

After being seated and fitted with the EEG cap, participants completed a short 1-minute familiarization task. During this task, the two reference images were presented successively 10 times each, for 2500 ms per image (inter-stimulus interval [ISI] = 500 ms). Each image was paired with either a 200-ms low-pitched (250 Hz) or high-pitched (290 Hz) sound, counterbalanced across participants. Participants were instructed to memorize the visual features of each picture (e.g., size, shape, colour) and the corresponding sound. At the end of this phase, the experimenter then conducted a control check by asking participants to recall which sound was associated with each image. All participants provided correct answers at this stage, suggesting that they successfully learned the two sound-image pairs.

All participants completed the two tasks in the same fixed order, with the imagery task first to minimize participants’ fatigue during this task. Both tasks were designed to be as similar as possible (Figure 2) and were preceded by a small 2-trial training phase with each reference image (cat and strawberry) presented once. At the end of each training, the experimenter came back in the testing room and asked participants to explain, in their own words, the instructions for the subsequent task to ensure good comprehension and clarify any misunderstanding.

**Figure 2.**
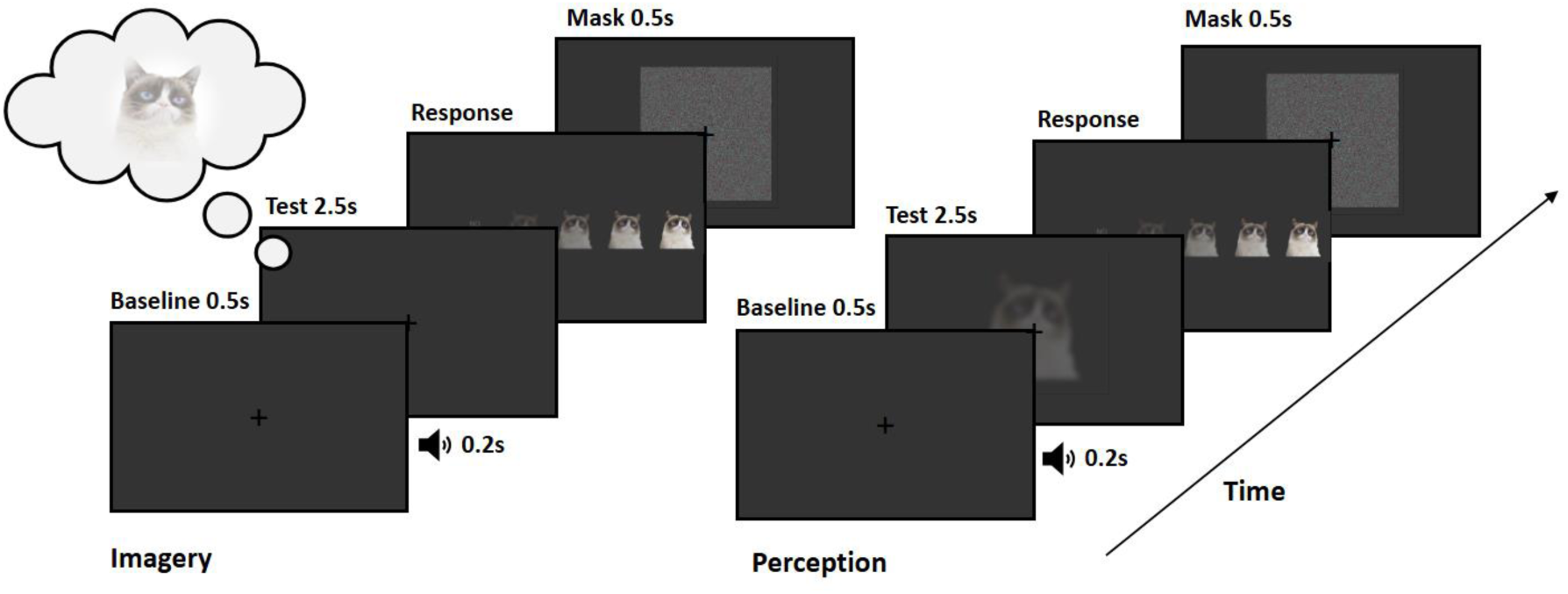
Illustration of the trial sequence during the imagery and perception tasks.

Within each task, every trial began with a grey 500-ms baseline screen displaying a fixation cross, followed by a sound cue. The onset of the cue was randomized between 400 ms and 600 ms (in 20-ms steps) to minimise anticipation effects. The sound prompted participants to either imagine (in the imagery task) or view (in the perception task) the cued image for 2500 ms while the fixation cross remained on screen. Participants then selected the image that best matched how they imagined (or perceived) the cued stimulus from a set of five options using a mouse (see Supplementary Materials for all instructions). To prevent afterimages and clear the visual buffer before the onset of the next trial, a 500-ms dynamic-noise mask was presented immediately after each response.

Each task consisted of 300 randomized trials, with each category (cat or strawberry) cued 150 times. For the perception task only, each category was further divided into five vividness levels (coded from 1 = no image, to 5 = original image), each presented 30 times. Participants could take a self-paced break every 60 trials. The experiment typically lasted approximately 45 minutes.

### 2.5. Data preprocessing

Preprocessing choices can affect statistical significance and decoding accuracy, and practices are heterogeneous (Kessler et al., 2025; Widmann et al., 2015; Gabard-Durnam et al., 2018; Delorme, 2023). The preprocessing pipeline ensured the retention of meaningful data while avoiding overcleaning following recent recommendations.

For each participant, bad channels were identified and handled with the RANSAC algorithm, using the Python implementation of the Prep Pipeline (Appelhoff et al., 2025; Bigdely-Shamlo et al., 2015). All EEG artifacts were detected and reconstructed using Independent Component Analysis (ICA) with the Preconditioned ICA for Real Data (PICARD) algorithm (Ablin et al., 2018). To this aim, components corresponding to non-brain artifacts (e.g., eye, muscle, line noise) were automatically identified with the MNE-Python implementation of ICLabel (Pion-Tonachini et al., 2019). After cubic spline interpolation of any rejected channels based on the remaining channels, data was then filtered using a zero-phase FIR filter with a bandwidth between 0.1 Hz (as recommended by Widmann et al., 2015) and 42 Hz. The 42 Hz threshold for the low-pass filter was determined following MNE-Python recommendations and Nyquist theorem to avoid aliasing effects, as the signal had to be subsampled later i.e., one third of the desired frequency (128/3).

Relevant data segments were then epoched and baseline-corrected by subtracting the mean signal between -200 ms to 0 ms before event onset from the entire epoch. Metadata including task condition, stimulus category, vividness, and trial index were attached to each epoch.

### 2.6. Model design

#### 2.6.1 Input

To meet the model’s input requirements and reduce computational cost, epoched data were resampled to 128 Hz and cropped to a 0–1 s post-stimulus time window. The network therefore received single-trial EEG time series from 8 posterior electrodes (O1, O2, Oz, PO3, PO4, PO7, PO8, POz) across this time window (8 channels × 128 samples per trial).

The selected time window (0 – 1000 ms) aligned with previous studies showing that relevant visual processing steps occur within the first 1000 ms after stimulus onset (Carlson et al., 2013) and the claim that the typical duration of a mental image is as brief as 250 ms without active maintenance (Kosslyn, 1996). The 8 posterior electrodes were selected based on their proximity with EVAs (see Introduction). Supplementary control analyses further confirmed that this region of interest was the most relevant to vividness classification during the perception task (see Figures S1-S3 in Supplementary Analyses). No explicit time–frequency decomposition or band-specific features were extracted on the input data, as the frequency-sensitive representations that are the most relevant to the task are learned directly via temporal convolutional filters within the convolutional neural network (see next section).

#### 2.6.2. Architecture and parameters

To classify vividness levels, we used EEGNet, a compact convolutional neural network (CNN) initially designed for Brain-Computer Interface (BCI) applications, that is suitable for small EEG datasets while producing neurophysiologically interpretable features (Lawhern et al., 2018). The model was implemented using the reference architecture and parameters provided by the authors (https://github.com/vlawhern/arl-eegmodels).

Each input was passed through two sequential blocks of convolutional layers, followed by a classification layer. In the first block, EEGNet performed a temporal convolution (kernel length = 64 samples) to learn frequency filters and output eight frequency feature maps (F1 = 8). Through a depthwise convolution connected to each feature map individually, the network then learned two spatial filters (D = 2) per feature map, resulting in sixteen (F2 = 16) frequency-specific spatial filters. The second block consisted of a separable convolution that combined depthwise convolution followed by a pointwise convolution. The former learned a temporal summary for each feature map individually and the latter learned how to optimally mix the feature maps together. Finally, in the classification block, the extracted features were flattened and passed through a fully connected (‘dense’) layer to produce a vector containing raw scores for each class. A final SoftMax activation layer then converted the vector to a probability distribution over all possible classes. To regularize the model’s learning and help promote independence between feature maps learned by the model, two 2D-spatial dropout layers were employed at the end of each block, with a dropout probability set to 0.4. Full details of EEGNet architecture can be found in the reference article (Lawhern et al., 2018).

#### 2.6.3. Output

For each input, EEGNet outputs a probability distribution across the classes on which the model was trained. We computed two main variables from each outputted probability distribution: (1) a discrete predicted value, corresponding to the class with the higher probability, and (2) a continuous-valued estimate, referred to as ‘expected vividness’, which was computed as:

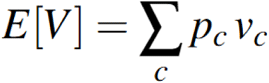

Where *p_c_* is the predicted probability for class *c*, and *v_c_* is the corresponding vividness level. Discrete predictions allowed us to monitor the model’s training and validation accuracy, and expected vividness allowed us to quantify vividness while preserving uncertainty and graded similarity between labels, therefore making intermediate predictions interpretable. This is particularly useful to compare vividness predictions in cases where tested trials did not have ‘true’ labels (e.g., imagery trials) and to assess how the model interpolated trained labels to predict unseen labels.

### 2.7. Training and testing logic

Classification was performed using a Leave-One-Subject-Out (LOSO) cross-validation approach i.e., all but one participant’s data was used to train the model while the held-out participant was used for testing, repeated until all participants have been tested (see Figure 3). The model was therefore trained 34 times (*N* = 34 participants) for 150 iterations with a minibatch size of 32 trials using the Adam optimizer (learning rate = 5e10-4, clip norm = 1.0) to minimize categorical cross-entropy loss. Predictions on unseen perception and imagery trials were embedded within the same LOSO procedure, to ensure that all predictions were strictly out-of-sample. Empirical chance levels were estimated via label permutations i.e., randomly shuffling the labels 100 times and testing the model predictions for each random permutation (Pereira et al., 2009).

**Figure 3.**
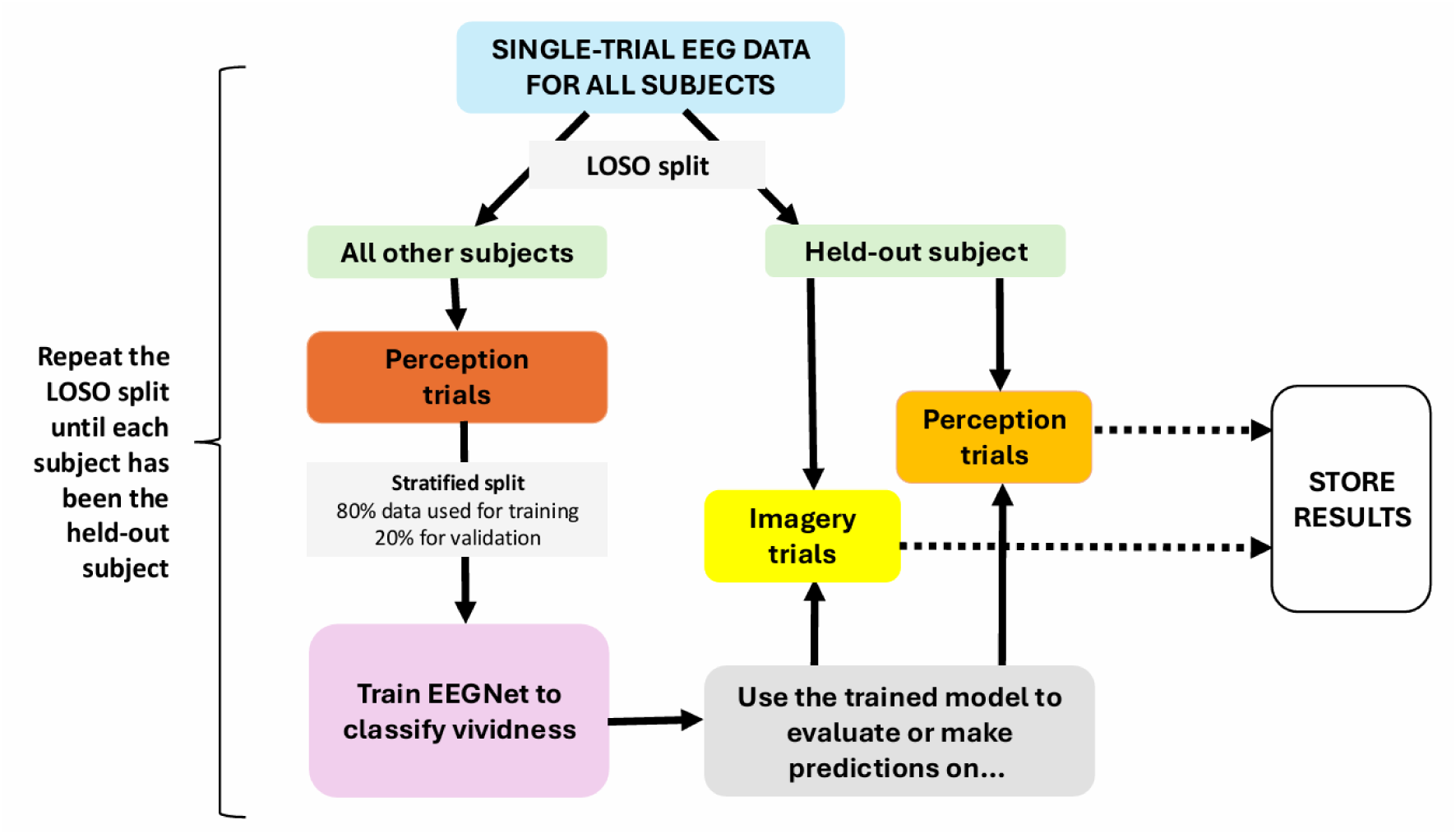
Training, validation, and testing procedure using leave-one-subject-out (LOSO) cross-validation.

### 2.8. Performance metrics

To evaluate performances on the vividness-classification task (balanced classes), we used three performance metrics: accuracy, F1-score and Quadratic-Weighted Cohen’s kappa (QWK), as implemented in sklearn (v1.7.1). Accuracy corresponds to the proportion of correctly reported or predicted labels relative to the total number of responses, reported as percentages in the manuscript. The F1-score uses the harmonic mean between Precision and Recall to provide a balanced index of both metrics (see Grandini et al., 2020). The QWK coefficient evaluates agreement between true and predicted or reported labels (Artstein & Poesio, 2008). Quadratic weighting was preferred over linear weighting to account for the fact that vividness levels may be ordinal, but their labels remain arbitrary and may not correspond to evenly spaced observations at the neural level.

### 2.9. Statistical analyses

We compared group-level differences in expected vividness between conditions or differences to chance using paired-samples t-tests (pingouin, version 0.5.5) with assumptions of normality and homoscedasticity assessed via Shapiro-Wilk and Levene’s tests (SciPy, version 1.16.1), respectively. To examine how expected vividness was affected by reported vividness at the trial level, we conducted mixed-effects linear regression using statsmodels (version 0.14.5), with models specified as expected_vivid ∼ reported_vivid + (1 + participant), and reported vividness as a categorical predictor. At the individual level, we assessed associations between mean scores using Spearman’s rank correlations (total and partial) as implemented in the pingouin package (version 0.5.5). Correlation coefficients were interpreted following conventional guidelines (ρ < .19 = “very weak”, .20–.39 = “weak”, .40–.59 = “moderate”, .60–.79 = “strong”, > .80 = “very strong”), with significance set at α = 0.05.

## 3. Results

### 3.1. Human classification

In the imagery task, the mean reported vividness was 3.71 (*SD* = 1.0, range 1-5). Looking at per-subject reported vividness, we could identify three participants (orange, pink and green dots; Figure 4) with a mean reported vividness that was strictly below 2 (with 1 = no image at all). Of all the participants, only these reported that their mental images of the stimuli more closely matched seeing the stimuli shown at the lowest level of vividness or below for more than 75% of their respective 300 trials during the imagery task. Such systematically low scores may reflect task disengagement, atypical cognitive profile or reduced overall imagery ability (i.e., mild or severe aphantasia; Zeman, 2024; Zeman et al., 2020). These participants were retained in subsequent analyses to preserve the full variability of imagery abilities in the sample; however, their data are highlighted in figures for transparency.

**Figure 4.**
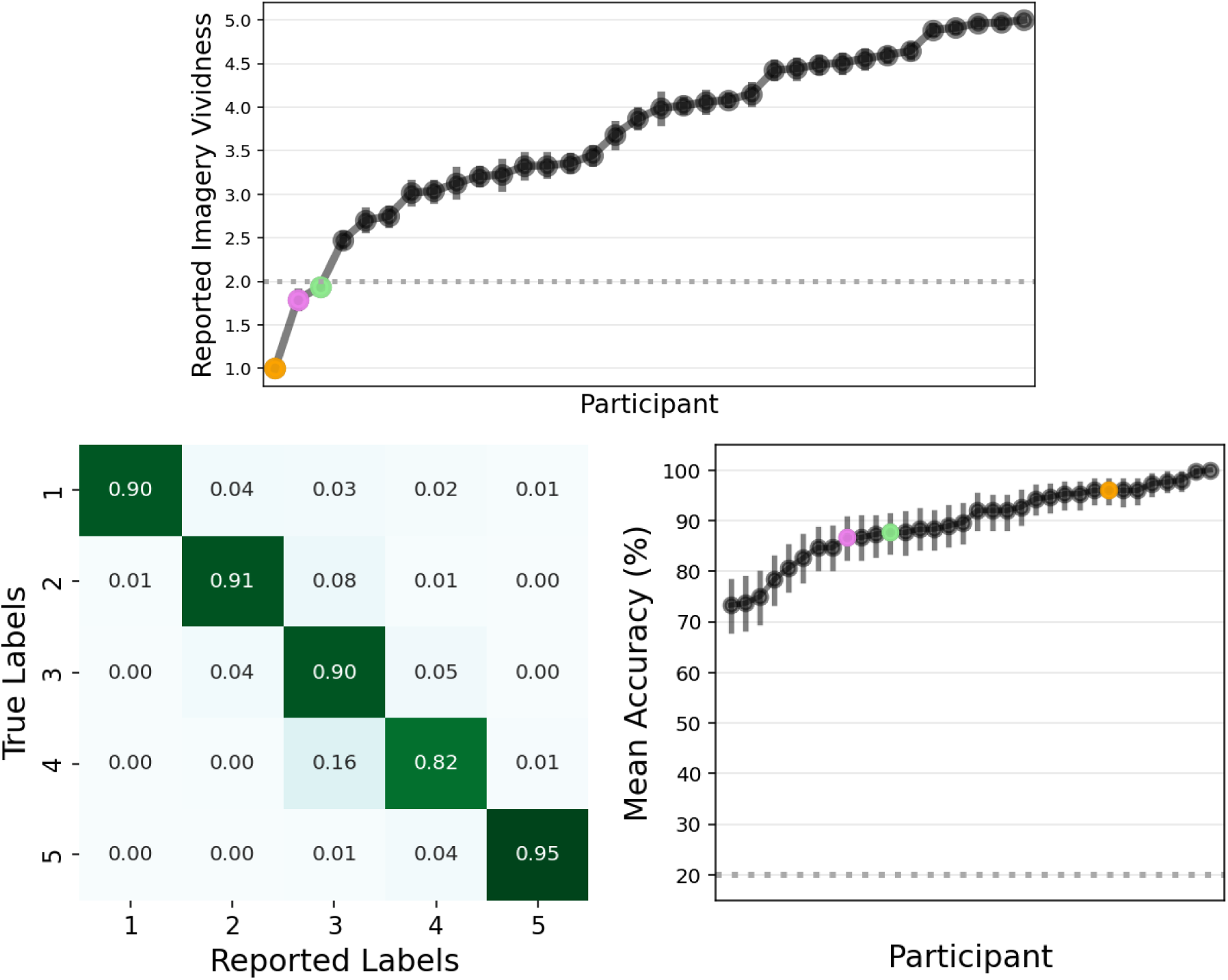
Confusion matrix showing 5-class human classification results, and individual variability in perception classification accuracy and reported imagery vividness. The coloured dots (pink, blue and orange) highlight the three participants with a mean reported VMI vividness strictly below two.

During the perception task, the mean accuracy across all five vividness levels per participant was 89.69% (*SD* = 7.34; range 73–100; F1-score: *M* = 0.90, *SD* = 0.08, range: 0.67-1.0) with no clear outlier. The mean reported vividness was 3.02 (*SD* = 0.09, range 2.90-3.37; true mean: 3.0). Agreement between participants’ reported vividness and true vividness labels was almost perfect (QWK = 0.94, *SD* = 0.06, range: 0.7-1.0).

### 3.2. Machine classification

#### 3.2.1 Five classes

##### 3.2.1.1. Performances

The EEGNet model was first trained to decode all five vividness levels seen during the perception task across all participants, on a predefined subset of eight posterior electrodes (see Methods section). The LOSO cross-validation procedure yielded a mean accuracy of 44.32% (SD = 6.18%), which was way above the 20% chance level (t(33) = 23.21, *p* < 0.001, *d* = 3.98, BF_10_ = 1.40 × 10^19^, *n* = 34). Despite noticeable individual variability (ranging from 31.67% to 55%), every single participant showed an above-chance mean classification accuracy, with no clear outlier (Figure 5). The mean F1-score suggested fair balance between Precision and Recall (*M* = 0.42, *SD* = 0.07, ranging 0.28 to 0.55), with moderate agreement between true and predicted labels (QWK: *M* = 0.52, *SD* = 0.14, ranging 0.25 to 0.77).

**Figure 5.**
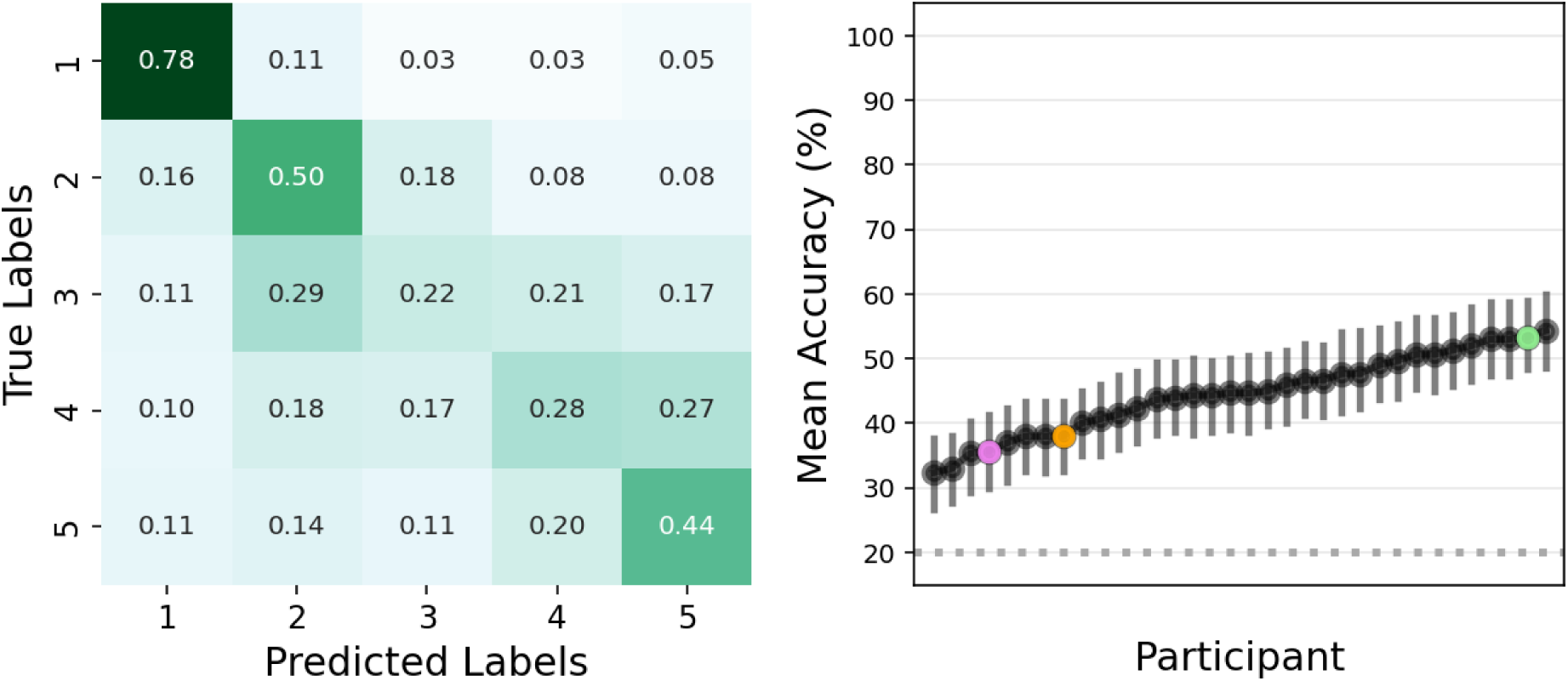
Confusion matrix for the 5-class EEGNet model and per-participant mean accuracy with 95% confidence intervals. Most misclassifications occurred between adjacent vividness levels, while levels 1, 2, and 5 were predicted more accurately. All participants had a mean accuracy above the chance level (dotted line; 20%). The coloured dots (pink, blue and orange) highlight the three participants with a mean reported VMI vividness strictly below two.

Further inspection of the 5-class confusion matrix (Figure 5) revealed that misclassifications were largely confined to cells adjacent to the diagonal. Notably, stimuli shown at intermediate vividness (level 3) exhibited substantial bidirectional confusion with neighbouring vividness levels; while the baseline screen, stimuli shown at the lowest vividness level and the original stimuli (i.e., levels 1, 2, and 5 respectively) were classified with higher accuracy. The fact that misclassifications were not uniform suggest that the model learned an ordinal or graded relationship between labels.

##### 3.2.1.2. Mean Expected Vividness Comparisons Between Classes

At the group level, we found extreme evidence that mean expected vividness (i.e., expectation over probability distribution; see Methods) while seeing the stimuli shown at the lowest vividness level (class 2) was greater than the mean expected vividness while looking at the baseline screen (class 1; t(33) = 25.05, *p <* 0.001, *d* = 3.81, BF_10_ = 1.40 × 10^20^) while remaining lower than mean expected vividness for stimuli shown at the intermediate vividness level (class 3; t(33) = 12.03, *p <* 0.001, *d* = 1.24, BF_10_ = 1.15 × 10^11^, *n =* 34). Mean expected vividness while seeing stimuli at high vividness (class 4) was greater than for this intermediate level (t(33) = 9.83, *p <* 0.001, *d* = 0.64, BF_10_ = 7.56 × 10^8^) but lower than for the original stimuli shown at the highest vividness level (class 5; t(33) = 6.46, *p <* 0.001, *d* = 0.34, BF_10_ = 1.28 × 10^5^, *n =* 34). At the individual level, this corresponded to 100.0% (34/34), 97.06% (33/34), 97.06% (33/34) and 85.29% (29/34) of participants showing a difference in the hypothesized direction for each between-class comparison, respectively. This result therefore strongly supports that, despite modest classification performances when considering discrete predicted values, expected vividness reliably preserved the ranks of the labels.

#### 3.2.2. Three classes

##### 3.2.2.1. Performances

To provide meaningful predictions of vividness on imagery trials, the model must achieve sensitive classification performances during VP but also be able to predict new vividness labels on which it was not trained with enough reliability. To improve classification performances and enable testing on new labels, a second EEGNet model was trained to classify only stimuli shown at three out of five vividness levels (Figure 6). These levels corresponded to labels 1, 2 and 5, previously identified as being the three most reliably separable vividness levels by the 5-class EEGNet (cf. Figure 5). Perception trials pertaining to labels 3 and 4 were set aside for downstream analyses.

**Figure 6.**
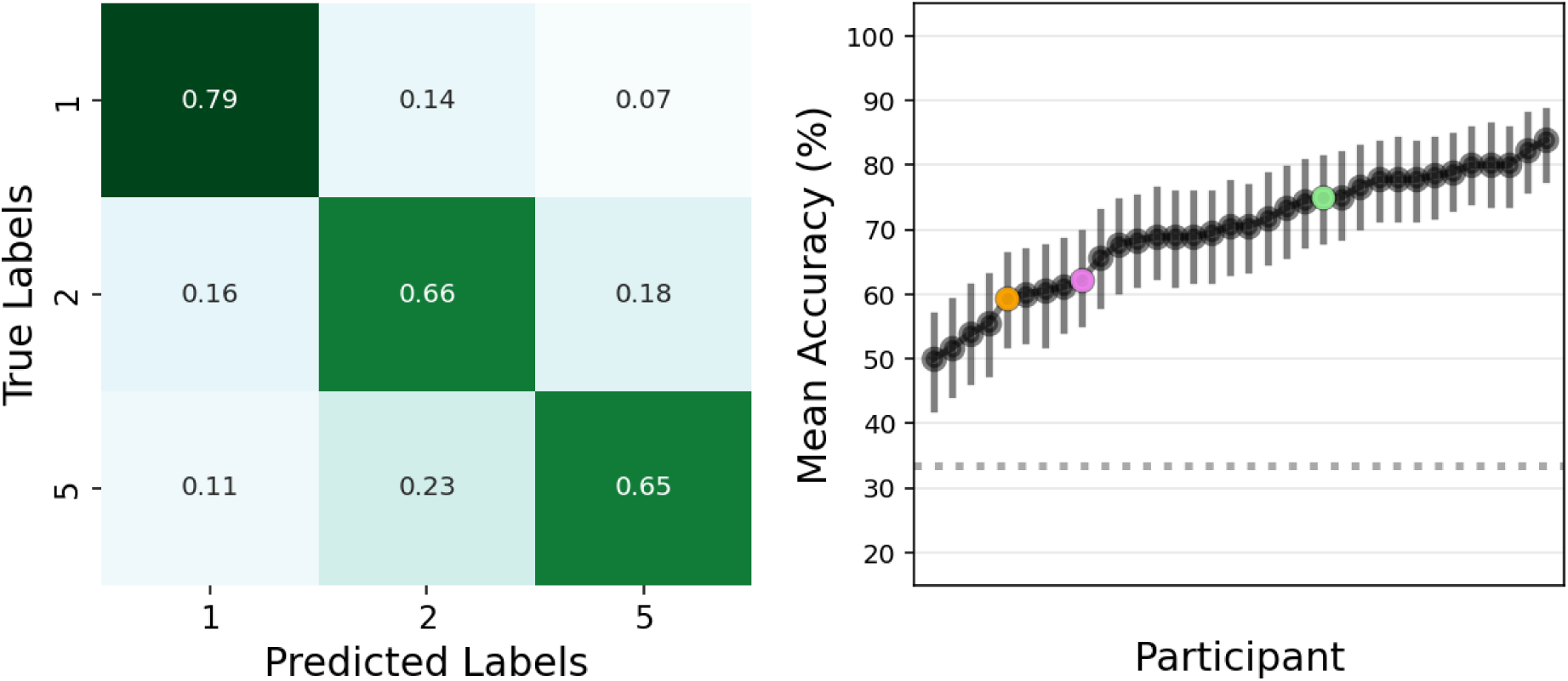
Confusion matrix for the 3-class EEGNet model and per-participant accuracy with 95% confidence intervals. Misclassifications were stronger between adjacent vividness levels. All participants had a mean accuracy above the chance level (dotted line; 33%). The coloured dots (pink, blue and orange) highlight the three participants with a mean reported VMI vividness strictly below two.

The LOSO procedure revealed a mean accuracy of 69.31% (*SD* = 8.09%, range 52.78–85.00%), which was also way above the 33% chance level (t(33) = 23.24, *p* < 0.001, *d* = 3.99, BF_10_ = 1.44 × 10^19^; *n* = 34). The mean F1-score increased accordingly to 0.69 (*SD* = 0.10, ranging 0.48 to 0.84), with a mean QWK that reflected substantial agreement between true and predicted labels (*M* = 0.63, *SD* = 0.14, ranging 0.35 to 0.83). While this simplification of the classification problem may not have significantly increased mean validation accuracy relative to chance, it reduced absolute misclassifications while reducing model complexity by 7.94% (5-classes: n(params) = 1637; 3-classes: n(params) = 1507).

##### 3.2.2.2. Mean Expected Vividness Comparisons Between Classes

Similar to the 5-class EEGNet, predictions with the 3-class EEGNet yielded a mean expected vividness that was significantly greater when participants viewed lowest-vivid stimuli (class 2) than during the baseline screen (class 1; t(33) = 20.16, *p* < 0.001, *d* = 3.21, BF_10_ = 2.10 × 10^17^, *n* = 34) but lower than mean expected vividness when participants were looking at the original stimuli (class 5; t(33) = 12.64, *p* < 0.001, *d* = 2.12, BF_10_ = 4.17 × 10^11^, *n* = 34, *n* = 34). This corresponded to 100% (34/34) and 97.06% (33/34) of participants showing a difference in the hypothesized direction, respectively.

##### 3.2.2.3. Generalization to Unseen Perception Trials

The main advantage in having a model trained on 3 out of 5 labels is that we can assess generalization to stimuli associated with the unseen vividness labels. To this aim, the trained 3-class EEGNet was then applied to the held-out participants’ remaining perception trials, while viewing stimuli at intermediate and high vividness levels (i.e., labelled as class 3 and 4). All predictions were embedded within the aforementioned LOSO procedure i.e., generated fully out-of-sample at the participant level, such that no neural data from the test participant could have contributed to model training (see Methods section).

Prediction results on the unseen perception trials confirmed that expected vividness tracked the ordinal nature of the actual vividness labels. At the individual level, participants had a mean expected vividness for label 4 (*M* = 3.35, *SD* = 0.49) that was higher than for label 3 (*M* = 3.05, *SD* = 0.42; t(33) = 8.34, *p <* 0.001, *d* = 0.64, BF_10_ = 1.84 × 10^7^, *n* = 34). Mean expected vividness for label 3 was significantly greater than for class 2 (t(33) = 11.30, *p* < 0.001, *d* = 1.22, BF_10_ = 2.29 × 10^10^), and label 5 significantly greater than label 4 (t(33) = 4.46, *p* < 0.001, *d* = 0.36, BF_10_ = 555, *n* = 34). More specifically, this corresponded to 91.18% of all participants (31/34) with a mean expected vividness greater for class 4 than class 3, 97.06% with class 3 greater than class 2 (33/34), and 85.29% (29/34) with class 5 greater than class 4.

To summarize, we demonstrated that a compact CNN model could decode the vividness of perceived stimuli above chance and generalize across participants. Successful interpolation between trained categories confirmed that EEGNet learned a graded neural signature of vividness rather than merely memorizing categorical labels, and ensured the sensitivity of expected VP vividness (neural level) to true variations in stimulus-level vividness. Supplementary saliency analyses further suggested that both models relied on meaningful spatiotemporal patterns rather than gross amplitude differences (see Figures S4-S8 in Supplementary Analyses). Overall, these findings support a shared and generalizable neural signature of VP vividness across participants.

### 3.3. Generalization to Imagery Trials and Comparisons with Perception Trials

#### 3.3.1 Five classes

Embedding imagery testing into the LOSO loop (see Methods section), predictions on the unseen imagery trials yielded a mean expected vividness of 2.12 (SD = 0.18, range 1.77–2.46) during the imagery task. In the current framework however, expected imagery vividness can only be understood and interpreted in comparison to expected vividness during the perception task (Figure 7).

**Figure 7.**
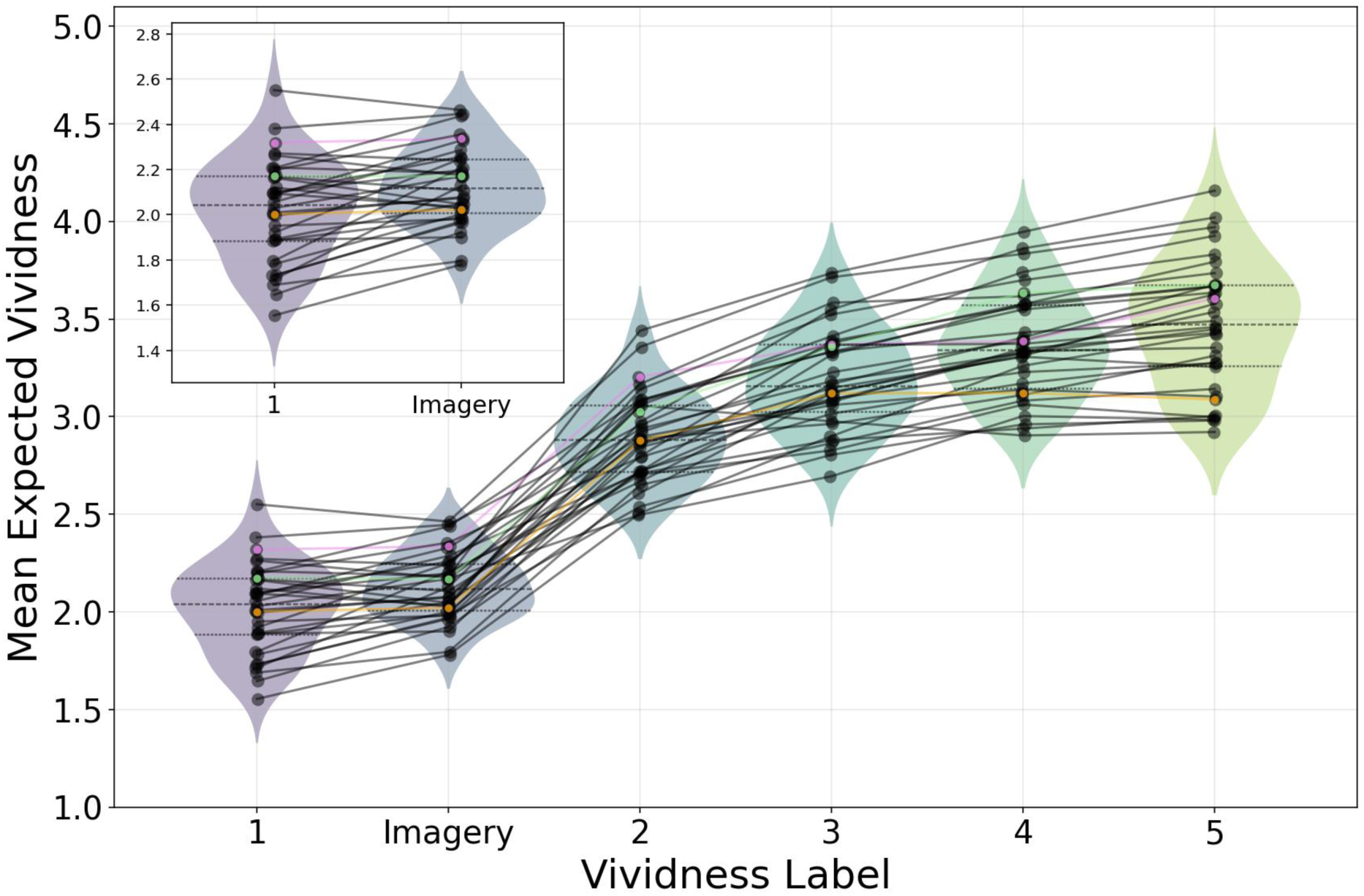
Mean expected vividness across classes during perception and imagery for the 5-class EEGNet, for each participant. Violin plots depict the group-level distribution across participants, with embedded quartiles; black dots indicate individual participant means. Thin black lines connect within-subject estimates across conditions. Coloured dots (pink, blue and orange) highlight the three participants with a mean reported VMI vividness strictly below two.

At the group level, we found that mean expected vividness during the imagery task was significantly greater than mean expected vividness for the baseline screen (class 1) during the perception task, supported by extreme evidence towards the existence of a true difference (t(33) = 4.38, *p* < 0.001, *d* = 0.53, BF_10_ = 456) but also remained lower than mean expected vividness for the stimuli shown at the lowest vividness level (class 2: t(33) = 20.90, *p* < 0.001, *d* = 3.73, BF_10_ = 6.07 × 10^17^, *n =* 34). Specifically, this corresponded to 73.53% (25/34) and 100% (34/34) participants showing a difference in the hypothesized direction for the comparisons with class 1 and 2 respectively.

For each participant, we then expressed mean expected VMI vividness as a percentage change relative to the mean expected vividness for the original stimuli (class 5) or baseline screen (class 1), using the formulas *(expected – max)/max × 100* and *(expected – min)/min × 100*, respectively. Expected vividness during the imagery task was on average 38.26% lower than expected VP vividness for the original stimuli (*SD* = 6.85, range 50.57–23.86%), and 5.98% higher than expected perceptual vividness when nothing was shown on screen during the perception task (*SD* = 7.84, ranging -5.41 to 24.13%, *n =* 34).

#### 3.3.2 Three classes

Prediction results on the unseen imagery trials revealed a mean expected vividness of 1.88 (*SD* = 0.17, range 1.50–2.23) during the imagery task (Figure 8). Comparing expected vividness between VP and VMI at the group level, expected VMI vividness was also significantly greater than the mean expected VP vividness of class 1 (t(33) = 3.97, *p* < 0.001, *d* = 0.45, BF_10_ = 156), but lower than class 2 (t(33) = 17.20, *p* < 0.001, *d* = 3.01, BF_10_ = 2.06 × 10^15^, *n =* 34), both comparisons supported by extreme evidence towards the alternative hypothesis. More precisely, this once again corresponded to 73.5% (25/34) participants with a mean expected vividness greater during the imagery task than during the baseline screen (class 1) shown during the perception task; and 100% (34/34) participants with a mean expected vividness lower during the imagery task than when seeing stimuli at the lowest vividness level (class 2) during the perception task. Using the same ratio calculations as described previously, expected imagery vividness was on average 45.75% lower than expected perceptual vividness for the original stimuli (*SD* = 7.97, range 61.38–29.29%), and 5.62% higher than expected perceptual vividness when nothing was shown on screen during the perception task (*SD* = 8.10, range -8.4 to 23.77, *n =* 34).

**Figure 8.**
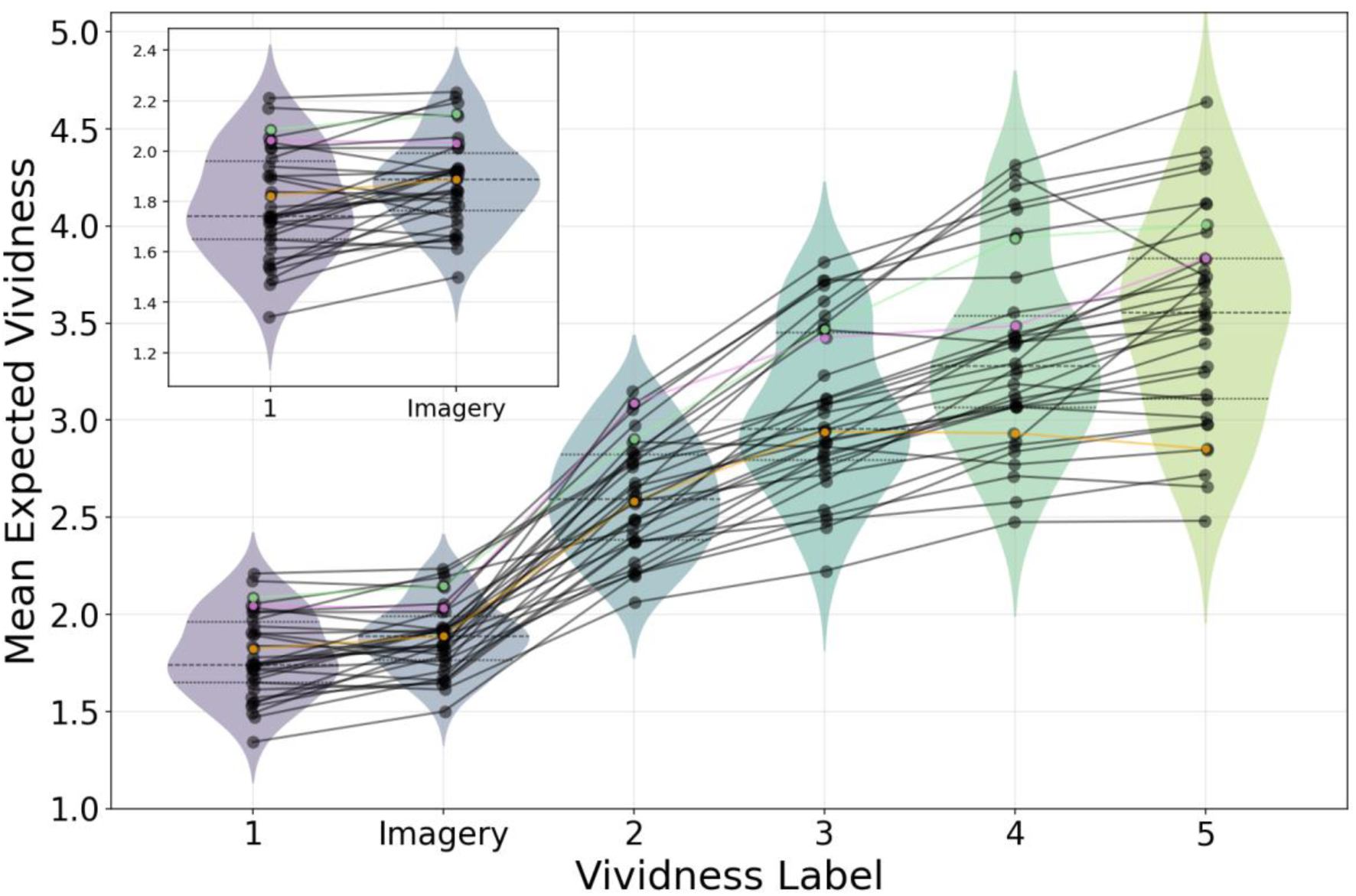
Mean expected vividness across classes during perception and imagery for the 3-class EEGNet, for each participant. Violin plots depict the group-level distribution across participants, with embedded quartiles; black dots indicate individual participant means. Thin black lines connect within-subject estimates across conditions. Coloured dots (pink, blue and orange) highlight the three participants with a mean reported VMI vividness strictly below two.

### 3.4. Relationship Between Mean Expected and Reported Imagery Vividness

#### 3.4.1 Five classes

We previously observed that during the imagery task, 31 out of 34 participants reported experiencing mental pictures that more closely matched seeing the actual stimuli at a vividness level ranging from low to as vivid as the reference image (see Human Classification results), suggesting a subjective estimation that was considerably higher than our neural-based estimation of VMI vividness relative to VP. To investigate whether expected VMI vividness in EVAs nonetheless scaled with phenomenological imagery vividness, we looked at the relationship between vividness as predicted by EEGNet and vividness as reported by participants during the imagery task (Figure 9).

**Figure 9.**
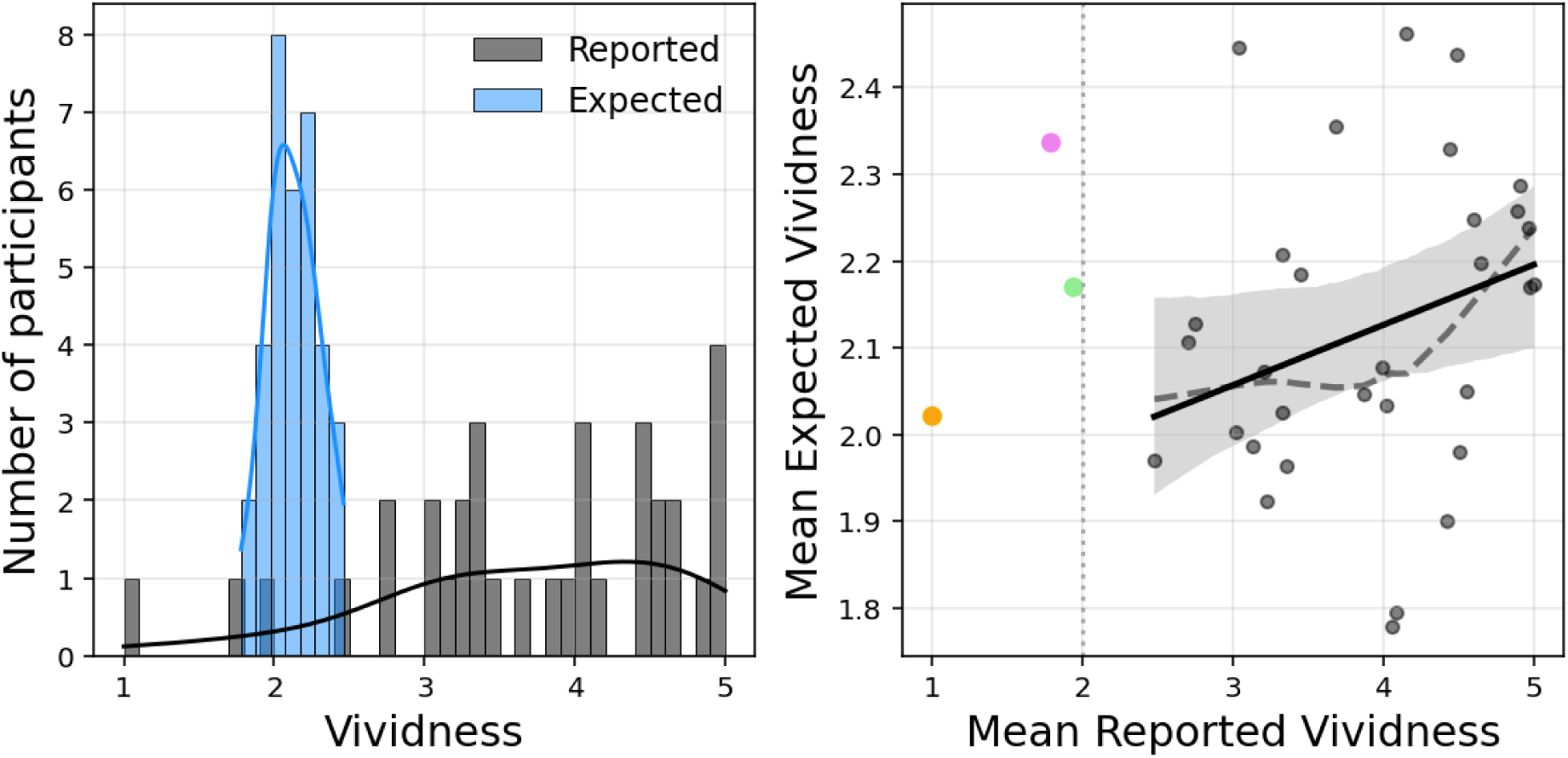
Relationship between reported and expected vividness for the 5-class EEGNet. Left panel: Distribution of mean self-reported vividness (black) and mean expected vividness (blue) across participants. Histograms are shown with kernel density estimates (bin width = 0.1), illustrating the central tendency and dispersion at the participant level. Right panel: Association between mean reported vividness and mean expected vividness across participants. Each point represents one participant; the solid line indicates the least-squares regression fit with 95% confidence interval shading and the dashed line the non-parametrical locally weighted linear regression fit (lowess). A vertical dotted line marks a report value of 2, used as a reference threshold for low reported vividness. The coloured dots (pink, blue and orange) highlight the three participants with a mean reported VMI vividness strictly below two.

At the trial level, although a subtle trend could be noticed, expected neural vividness was not significantly different between trials in which participants reported no image and those in which they reported vivid imagery (reported level 1 vs. reported levels 2-5: β = [0.013, 0.024, 0.032, 0.047], *SE* = [0.041, 0.039, 0.039, 0.047], *z* = [0.316, 0.615, 0.815, 1.143], *p* = [0.752, 0.539 0.415, 0.253]). At the individual level, the correlation between mean expected imagery vividness and mean reported vividness showed a positive non-significant relationship, with anecdotical evidence towards the null hypothesis that there is no relationship (Spearman’s correlation: ρ = 0.23, 95% CI = [-0.06, 1.00], *p* = 0.095, BF_10_ = 0.874, power = 0.375, *n* = 34).

Now from the behavioural results, we had earlier spotted three participants with a mean reported vividness strictly lower than 2 (suspected mild or total aphantasia; *n* = 3; see Human Classification results). These three participants were also the only participants to have a mean expected vividness during the imagery task that was greater than their mean reported vividness (see Figures S9 and S10 in Supplementary Analyses). To examine whether participants with systematically low-vividness responses attenuated the relationship, we tested the correlation again without these three participants. For the remaining 91% of the sample, the correlation was significant (ρ = 0.31, 95% CI = [0.01, 1.00], *p* = 0.042, BF_10_ = 1.76, power = 0.543, *n* = 31), showing a positive albeit weak correlation supported by anecdotical evidence towards the alternative hypothesis that there is a positive relationship. The correlation was still significant after controlling for both behavioural (human) and EEGNet’s accuracy during the perception task (Spearman’s partial correlation: ρ = 0.34, 95% CI = [0.03, 1.0], *p* = 0.036, *n* = 31).

#### 3.4.1 Three classes

Comparisons using expected vividness as predicted by the trained 3-class EEGNet (Figure 10) revealed that expected vividness once again did not differ between trials in which participants reported no image and trials in which they reported more vivid imagery (reported 2-5: β = [0.015, 0.013, 0.030, 0.040], *SE* = [0.040, 0.039, 0.039, 0.041], *z* = [0.365, 0.326, 0.771, 0.991], *p* = [0.715, 0.745, 0.441, 0.321]). At the individual level, the correlation between mean expected vividness and mean reported vividness was positive but also not significant (ρ = 0.20, 95% CI = [-0.09, 1.00], *p* = 0.124, BF_10_ = 0.706, power = 0.318, *n* = 34). Excluding outlier participants with systematically low reported imagery vividness (*n* = 3), we found moderate evidence supporting that mean expected vividness correlated positively albeit moderately with mean self-reported vividness (ρ = 0.40, 95% CI = [0.11, 1.00], *p* = 0.014, BF_10_ = 4.441, power = 0.727, *n* = 31). The correlation remained significant even after controlling for behavioural and model performance during the perception task (ρ(partial) = 0.42, 95% CI = [0.12, 1.0], *p* = 0.012, *n* = 31).

**Figure 10.**
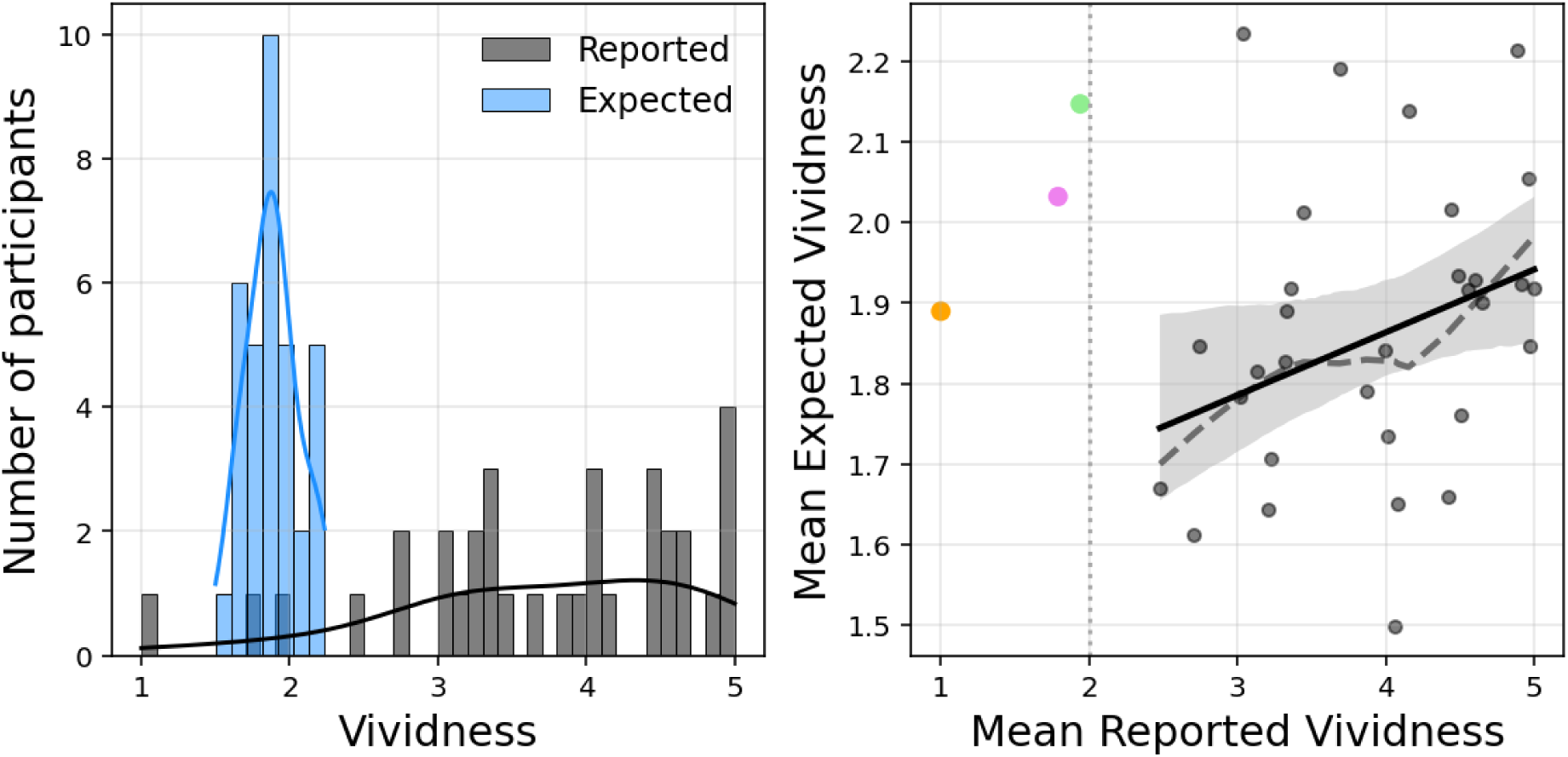
Relationship between reported and expected vividness for the 3-class EEGNet. Left panel: Distribution of mean self-reported vividness (black) and mean expected vividness (blue) across participants. Histograms are shown with kernel density estimates (bin width = 0.1), illustrating the central tendency and dispersion at the participant level. Right panel: Association between mean reported vividness and mean expected vividness across participants. Each point represents one participant; the solid line indicates the least-squares regression fit with 95% confidence interval shading and the dashed line the non-parametric locally weighted linear regression fit (lowess). A vertical dotted line marks a report value of 2, used as a reference threshold for low reported vividness. The coloured dots (pink, blue and orange) highlight the three participants with a mean reported VMI vividness strictly below two.

## 4. Discussion

In this study, we tested the feasibility of predicting vividness from single-trial EEG activity in early visual areas (EVAs) using a convolutional neural network, EEGNet. We found that EEGNet could learn a neural representation of vividness that generalized across participants and to unseen perception trials. When projecting imagery trials onto this perceptual neural space and evaluating the model’s predictions, mean expected imagery vividness was greater than expected vividness during the baseline screen but lower than while seeing stimuli shown at the lowest vividness level. For most participants, expected VMI vividness relative to VP scaled with their reported subjective experience of vividness.

The current methodological framework relied on four main assumptions: (1) vividness can be operationalized as variations in low-level features such as clarity and intensity, (2) neural representations in EVAs convey information about these low-level features, (3) VMI and VP share (at least some) of these neural representations in EVAs, and (4) EEG and brain decoding techniques can be used to retrieve this information from the neural signal (see Introduction). We successfully confirmed assumptions 1, 2 and 4 through above-chance classification accuracy and sensible predictions during the perception task; supporting the existence of a subject-independent, graded neural signature of vividness in EVAs when vividness is operationalized as a variation in low-level features such as clarity and intensity (Fazekas, 2024; Marks, 2023). This result echoes a recent study which re-analysed fMRI and MEG data in two perception tasks and similarly found evidence of a graded neural signature of perceptual vividness during VP (Barnett et al., 2024). Compared to the present study however, Barnett et al. (2024) employed self-reports of visibility and awareness (i.e., labelled vividness at the phenomenological level) and consequently found a neural signature of vividness that spanned not only visual areas, but also frontal and parietal areas. This suggests that the neural signature of vividness depends on how vividness is defined and measured, highlighting the importance of clear and explicit operationalisation.

The third assumption that VMI and VP share at least some neural representations in EVAs (Dijkstra, 2024; Huang et al., 2023a; Pearson & Kosslyn, 2015; Slotnick et al., 2005) remains debated (Bartolomeo et al., 2020; Spagna et al., 2021). In our approach, we relied on this assumption as a working framework to quantify imagery vividness relative to perception at the neural level. This was done by first establishing a perceptual vividness ‘scale’ based on how EVAs reacted to objective variations in low-level stimulus features during VP and then projecting VMI onto this ‘scale’ to estimate how imagery resembled perception at different levels of perceptual vividness. Although the spatial resolution of EEG makes it difficult to localise EVAs perfectly, we selected the electrodes that were most likely to capture the signals originating from the EVAs. Spatial saliency maps confirmed that this region of interest was the most relevant for distinguishing between levels of perceptual vividness. If VMI recruitment of EVAs were identical to EVAs recruitment while seeing the baseline screen, predictions along this perceptual scale would collapse to baseline; yet, we found extreme evidence that imagery predictions do fall slightly above the perceptual baseline while remaining well below high-vividness perception. This confirms that EVAs can contribute at least some weak, perceptual-like information during imagery. Importantly, our findings does not imply that perception and imagery share *identical* neural coding schemes but provide a relative quantification of imagery vividness along a perceptual reference. Within this framework, our findings support previous accounts that VMI is “quasi”, or, in this case, “barely” depictive (Kosslyn et al., 2006; Pearson, 2019; Pearson et al., 2015), in that early visual engagement is present during imagery but substantially attenuated compared to actual sensory input.

Interestingly, despite evidence of a VMI that is barely pictorial at the neural level, most participants still reported experiencing picture-like VMI during the imagery task. To explain this discrepancy, one could first argue that EEGNet’s vividness predictions simply cannot be trusted. Indeed, the reliability of VMI predictions also depends on the model’s ability to learn the relevant features—or in this case, echoing Naselaris et al. (2015), on how well-tuned the model is to low-level visual features—during VP. This corresponds to our fourth assumption (see Introduction). We believe this explanation to be quite unlikely, as we showed that EEGNet achieved a mean accuracy that was around two times above chance for both three and five classes classification, which is comparable to the mean VP accuracy achieved in other similar recent studies (e.g., Chang et al., 2025; Kilmarx et al., 2024; see Wilson et al., 2024 for a review). Based on this criteria, VMI predictions in the current study are therefore at least as reliable as VMI cross-decoding results in comparable studies. Moreover, the fact that between-class comparisons of expected vividness, a probabilistic index that preserved the model’s uncertainty, sensitively reflected vividness label ranks during VP for seen and unseen labels further supports the reliability of our measurement. Future studies aiming at improving classification accuracy during VP may nonetheless seek optimizing EEGNet training parameters via a grid search instead of using default parameters, refining the model architecture to better suit ordinal classification (e.g., Bérchez-Moreno et al., 2025, Lv et al., 2025) or using deeper networks (e.g., DeepConvNet: Schirrmeister et al., 2017) while keeping in mind that this comes at a cost of lower interpretability and greater risk of overfitting.

A more plausible interpretation of this discrepancy is that participants exploited the picture-based scale the same way they would for a Likert scale. Indeed, the use of picture-based reports allowed a consistent cross-task operationalization of vividness while eliminating many language-based biases that typically occur with a Likert-scale response format (Sulfaro, 2024). Yet the available range of response options may have implicitly forced participants to reinterpret the instructions: faced with a majority of pictorial response options, participants therefore decided to respond using a more absolute or ambiguous scale e.g., ‘I will select the picture with the highest vividness level because the mental image I had in my mind *felt* as vivid as real seeing’, or ‘because it *felt like* I could project my mental image on the screen as vividly as if it was there’ (Schwarzkopf, 2024). Similarly, we created vividness versions of the pictures by degrading and mixing its two main subdimensions: clarity and intensity (Fazekas, 2024; Marks, 2023). However, if clarity inherently plays a bigger role in how participants subjectively evaluate the vividness of their imagery (Huang et al., 2025), using pictures that mixed both subdimensions could also cause such discrepancy or lower-than-expected correlations. While the focus of the current study was on quantifying neural-level VMI relative to VP rather than to subjective VMI, further investigations with the use of an extended scale and greater control over the physical parameters of the stimuli could help clarify the respective role of clarity and intensity in both phenomenological and neural vividness.

Now it is also possible that VMI vividness in EVAs (neural level) may, in some cases, be independent of reported VMI (phenomenological level) or preconscious. For example, the neural signals in EVAs exploited by EEGNet to predict vividness could have indexed visual properties of the neural representations *before* active maintenance (Kosslyn, 1996; Pearson et al., 2001) or amplification (Liu, 2026), and that greater expected vividness in EVAs does not guarantee successful maintenance or amplification. This aligns with the temporal dynamics revealed by our supplementary saliency analyses, which suggested that EEGNet mostly relied on the first 500 ms of each trial, with brain activity within the 0-300 ms time window showing stronger relevance to vividness classification; while the typical duration of a mental image without active maintenance is around 250 ms (Kosslyn, 1996). Considering introspection as a form of signal detection (Morales, 2024), without active maintenance, poorly or non-amplificated signal in EVA may therefore be experienced as less vivid or imageless (Chang et al., 2025) or even remain nonconscious memory reinstatements (Slotnick & Schacter, 2006). This could also explain why we found some evidence that expected VMI vividness still weakly-to-moderately correlated with subjective reports once excluding outlier participants with low-to-absent self-reported vividness.

Speculating further, this also questions whether individuals reporting barely pictorial or no imagery at all can truly be characterized as showing an imagery ‘deficit’ *per se*. After all, the weaker VMI effects typically observed in VMI-VP comparisons at the neural, physiological and behavioural level suggest that reports of low-to-absent VMI vividness may in fact more closely reflect the ‘true’ perceptual qualities of mental images than the medium-to-high vividness levels reported by the rest of the population. In this sense, individuals reporting lower-to-absent VMI vividness with preserved performances on visual tasks (e.g., Knight et al., 2026), could possibly be reinterpreted as showing more *balanced* top-down influences reflected by a more down-to-earth interpretation and use of vividness scales (e.g., ‘I will select the lowest vividness level or ‘no image’ because I believe it is barely or not comparable at all to actual perceptual input’). In comparison, stronger influence of top-down activity in hyperphantasia and typical imagers (e.g., Milton et al., 2021) could reflect an *overuse* of prior knowledge to inspect and report the vivacity of VMI. Given our relatively small sample, we cannot yet draw firm conclusions from this discrepancy, which would require a dedicated study with a larger sample that comprises individuals across the whole imagery spectrum. More broadly, we consider that correlations with subjective reports should be treated as a tool to interpret and contextualize other measurements rather than as a validation target (Sulfaro, 2024). Future research should help clarify how expected VMI vividness relative to VP relates to other measurements of imagery and vividness instead. For example, because EVAs are the first cortical areas to receive sensory input from the retina, expected VMI vividness in EVAs might better explain individual differences in the pupillary light response to imagined light than reported vividness (Kay et al., 2022; Gardner et al., 2026; Vanbuckhave et al., 2026). Associations with visual hallucinations occurrence could also be explored (Panigutti et al., 2026).

Moving forward, another important point to discuss is whether expected VMI vividness captured spontaneous or involuntary mental images rather than voluntary imagery during the imagery task. We have several reasons to think that this interpretation is unlikely. First, we recall that both the VP and VMI tasks were designed to maximize similarity (*ceteris paribus*), except for the explicit instruction to actively visualise the picture associated with the sound during the VMI task—which, by definition, corresponds to voluntary imagery. Second, all participants underwent a training phase prior to the experiment to learn to associate beep cues with picture categories. While this ensured that participants could not start imagining the correct stimuli *before* hearing the cue, in the perception task however, expecting seeing a picture on screen very likely led to the generation of a weak, spontaneous mental image upon hearing the sound cue (Cabbai et al., 2024). This implies that any neural activity during the VP task possibly reflected a mixture of spontaneous imagery and perception (consistent with Dijkstra & Fleming, 2023). In this context, even if hearing the sound cue initially induced an involuntary mental image during the VMI task, these would also have been present during the VP task, and especially during the baseline screen. To achieve above-chance classification, EEGNet therefore likely learned to exploit a neural signature of VP vividness that was independent of spontaneous imagery or mere visual preparation, considered as non-relevant to vividness classification (as the sound cued the category, but not the vividness level). In this case, our finding of overall greater expected vividness during the imagery task than during the baseline screen can be interpreted as evidence of neural representations that are more perceptually vivid during voluntary imagery than during any other non-relevant process occurring while seeing the baseline screen.

Overall, the proposed methodological frameworks along with our findings suggest that it is possible to quantify ‘how much weaker’ VMI is relative to VP, but more fine-grained estimations could be obtained with a few methodological improvements. Indeed, if vividness has a graded neural signature, there theoretically remain many more perceptual experiences between levels 1 and 2 that the visual system can dissociate and, as our results and the literature suggest, at least one of these intermediate VP levels should be closer to VMI than the baseline screen. This could be investigated in future studies, by presenting more low-vividness perceptual experiences below level 2, possibly oscillating around the vividness threshold at which perception and imagery may become undistinguishable (Dijkstra & Fleming, 2023). More specific adjustments to the current experimental design will depend on the research objectives. For example, the current study focussed on vividness as a feature of the visual information as encoded in EVAs during VMI and VP (neural level; see Introduction). Putting a stronger emphasis on phenomenological vividness might help provide more insights into individual differences in reported imagery and their relationship to conscious access (Liu, 2026; Scholz et al., 2026). If the objective is to measure the absence of phenomenological VMI in aphantasia, then optimizing Sensitivity (Recall) in the lower classes is critical to ensure that the model does not miss ‘true’ low-vividness individuals or trials. Ultimately, the core of this approach, which involves projecting imagery onto a perceptual axis, is flexible enough to be applied to various other sensory domains (e.g., auditory, auditory-verbal, etc.; Dawes et al., 2024; Nanay, 2018) and measurement methods (e.g., pupillometry data), as long as a clear operationalisation of the decoded features (here, vividness) is provided.

To conclude, our findings strongly support the view that VMI is barely pictorial, and that individual differences in the vividness of visual information as encoded in EVAs during VMI remain only weakly-to-moderately associated with individual differences in experienced vividness, even when using picture-based reports. Our methodological framework provides a principled way to quantify and compare VMI and VP on a shared neural-behavioural scale, with implications for studying individual differences and aphantasia.

## CRediT authorship contribution statement

**CV:** Writing – original draft, Visualization, Software, Methodology, Investigation, Formal analysis, Data curation, Conceptualization. **GG:** Writing – review & editing, Validation, Supervision, Resources, Project administration.

## Ethics

All participants gave their informed written consent before participating in the study, which was in accordance with the APA Ethics Code (https://www.apa.org/ethics/). The data collection procedure respected the General Data Protection Regulation (GDPR), and the protocol had been reviewed and approved by the Faculty Research Ethics and Integrity Committee (Project ID: 5826) prior to any data collection.

## Data availability

All relevant materials, anonymised data and analysis scripts will be made publicly available on the OSF repository of the project: https://osf.io/fp25t/

## Supplementary Analyses

### 1. Full-scalp classification

In the current study, our region of interest (i.e., 8 posterior electrodes, the ones closest to early visual areas) was theory driven (see Introduction and Methods sections). To explore the spatial information EEGNet spontaneously exploits to classify vividness when information from the full scalp is available, we re-trained the 5-class model on all 64 electrodes. The model was trained and validated using a stratified split, with 80% of the data used for training and 20% for validation. We present the results through confusion matrices (i.e., predicted versus true labels and probability distribution across classes) and gradient-based spatiotemporal maps (see next section) to highlight what contributed most strongly to EEGNet’s decisions.

**Figure S1.**
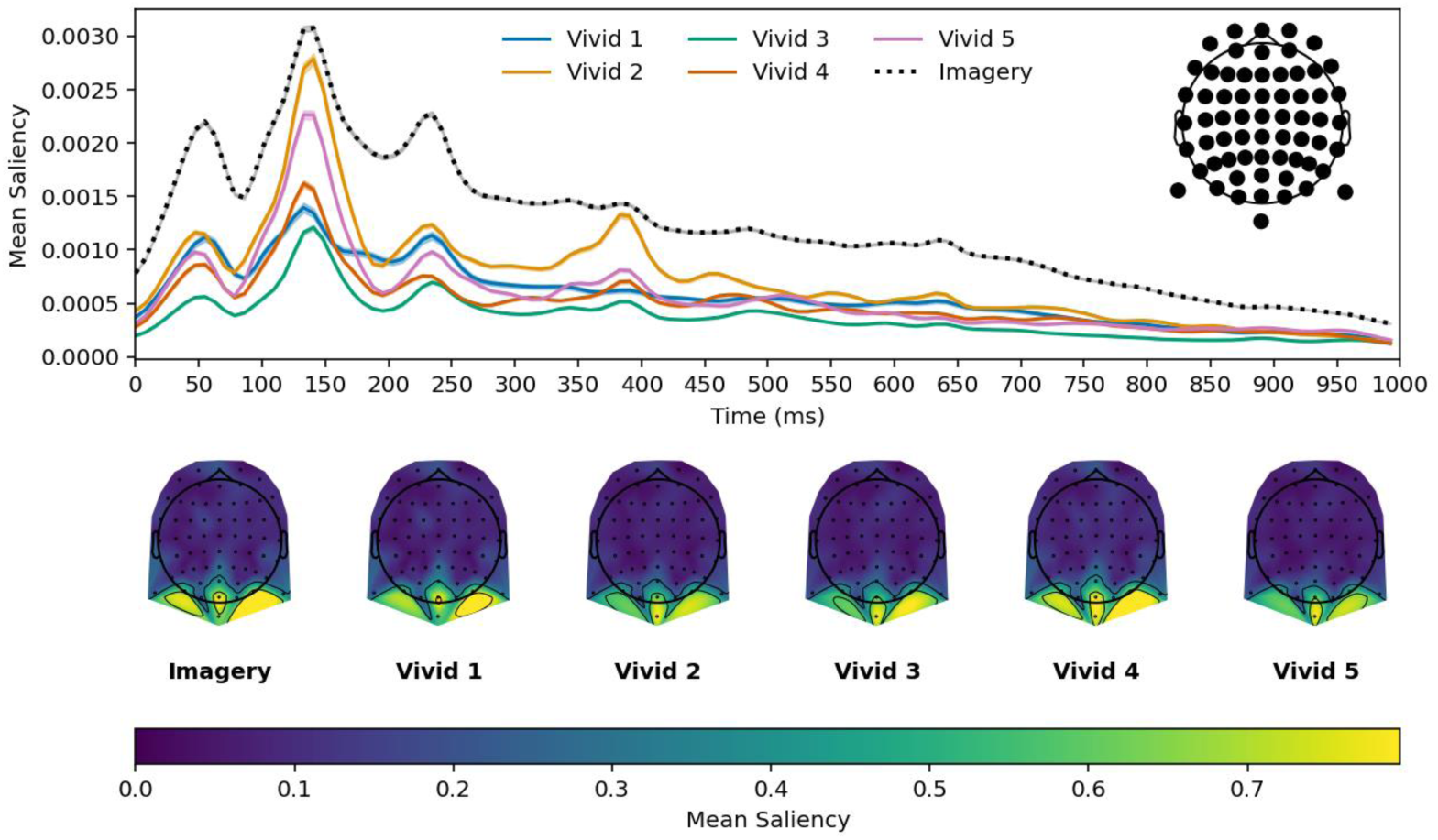
Mean temporal and spatial saliency maps for the 5-class EEGNet with 64 electrodes. This figure confirms that posterior electrodes contribute most to EEGNet’s predictions, consistent with visual cortex involvement in vividness encoding.

**Figure S2.**
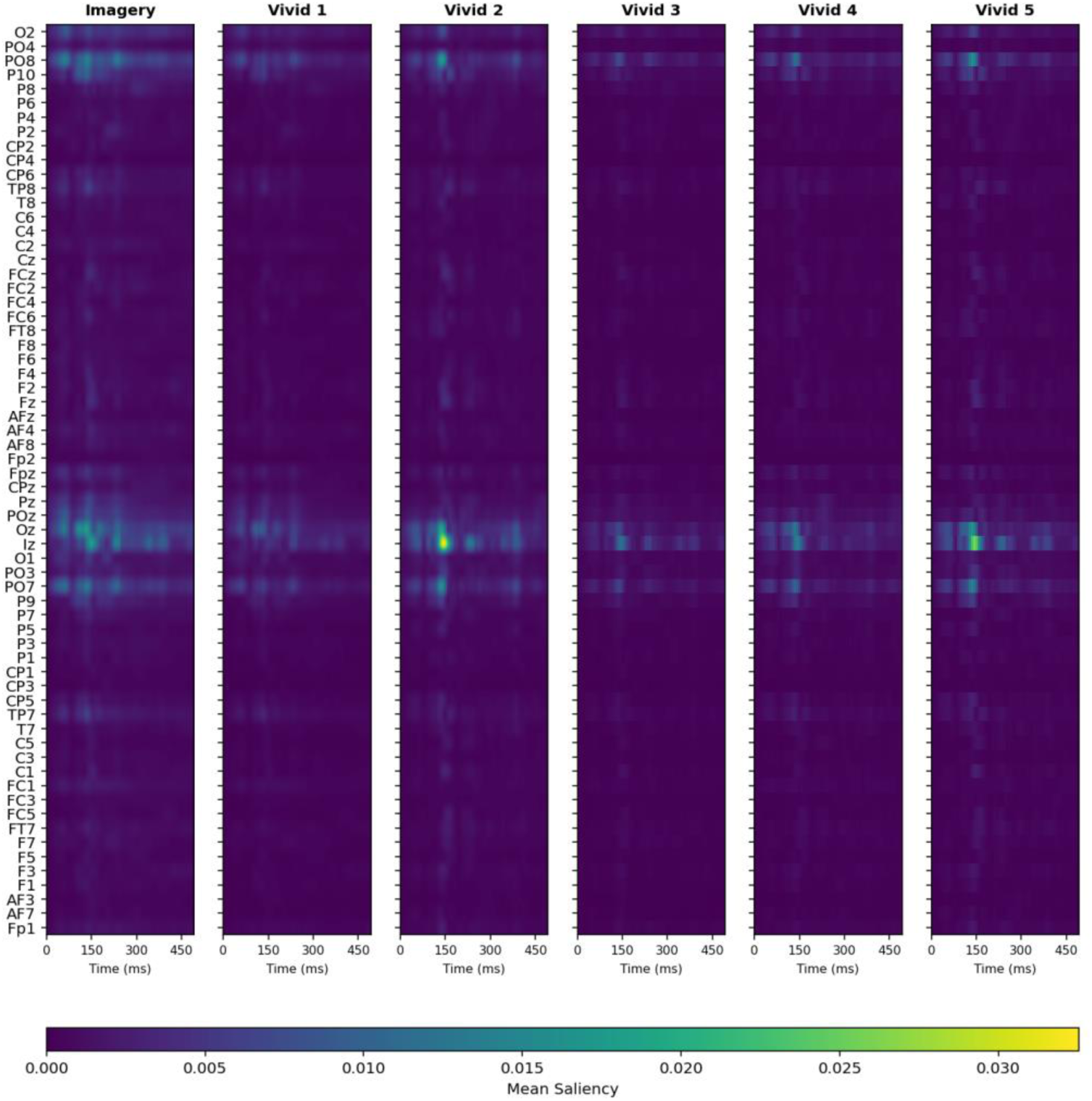
Spatial saliency maps per electrode during the first 500 ms after cue onset for the 5-class EEGNet trained on 64 electrodes. This figure confirms that posterior electrodes contribute most to EEGNet’s predictions, consistent with visual cortex involvement in vividness encoding.

**Figure S3.**
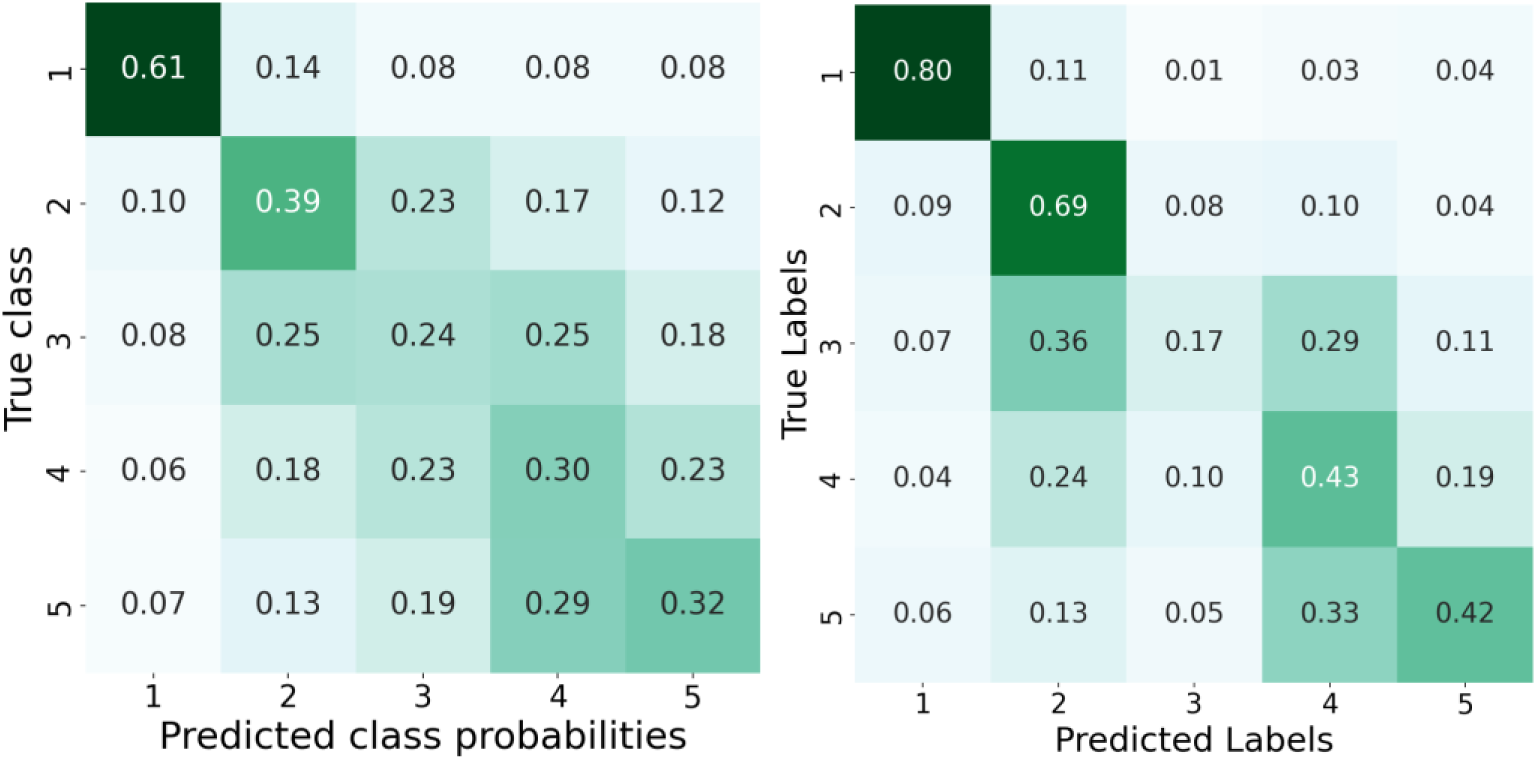
Confusion matrices of predicted labels and distribution of class probabilities for the 5-class EEGNet trained on 64 electrodes.

### 2. Gradient-Based Spatiotemporal Saliency Mapping

The EEGNet models reported in the main manuscripts were trained 34 times, on N-1 = 33 participants each time, and tested on the held-out participants (LOSO procedure). To compute general saliency maps in this section, we retrained the models on all N = 34 participants using a stratified split instead (no LOSO), with 80% of the data used for training and 20% for validation. Model performances were similar to the ones described in the main text with LOSO.

To investigate how EEGNet exploits spatiotemporal information in the EEG signal to discriminate between classes and predict imagery vividness, we employed a gradient-based saliency approach, which quantifies the contribution of each input feature to the model’s prediction by computing the gradient of the output with respect to the input EEG signal (Simonyan et al., 2014). Although more sophisticated attribution methods exist (Shrikumar et al., 2017; Sujatha Ravindran & Contreras-Vidal, 2023; Wang et al., 2023), the gradient-based method is still extensively used due to its computational simplicity and direct mapping from input features to output relevance (for a review, see: Ancona et al., 2018).

For each trial, gradients were computed using TensorFlow’s automatic differentiation function (GradientTape, v2.20.0) and saliency was calculated as the absolute value of the element-wise product of the gradient and the input signal (Grad × Input), yielding trial-wise relevance scores across all channels and time points, for each condition. This approach allows us to estimate how small perturbations in voltage at a given electrode and time point would influence the model’s evidence for that class, regardless of direction.

Temporal relevance was visualized by averaging saliency values across channels for each condition and plotting them over time. Spatial relevance was visualized by averaging saliency values across trials for each EEG channel, time point and condition. The resulting saliency maps therefore reflect the local sensitivity of the classifier’s decision to spatiotemporal features of the EEG signal, highlighting the time points and electrodes where changes in neural activity most strongly modulate the model’s output.

**Figure S4.**
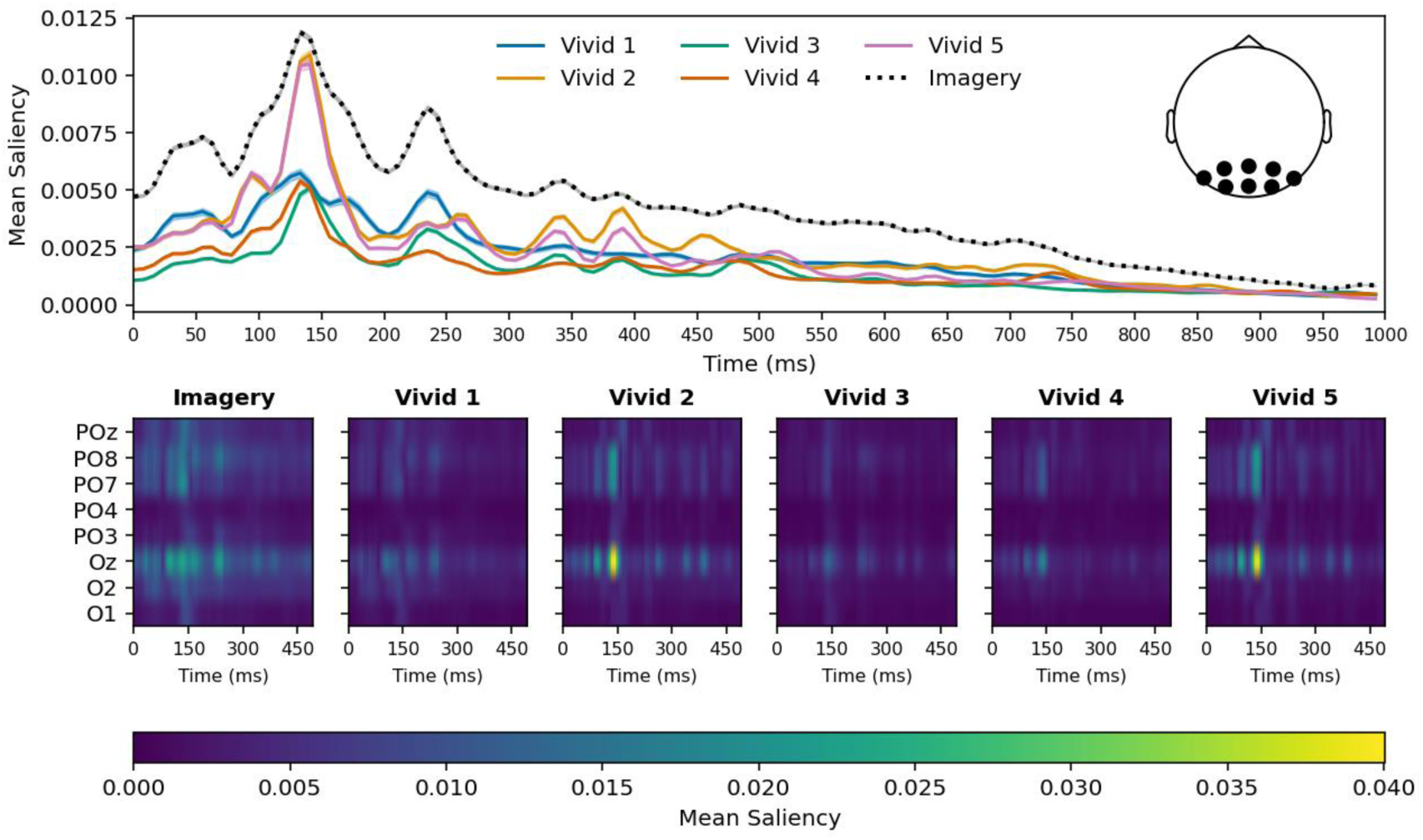
Mean temporal and spatial saliency maps for the 5-class EEGNet.

**Figure S5.**
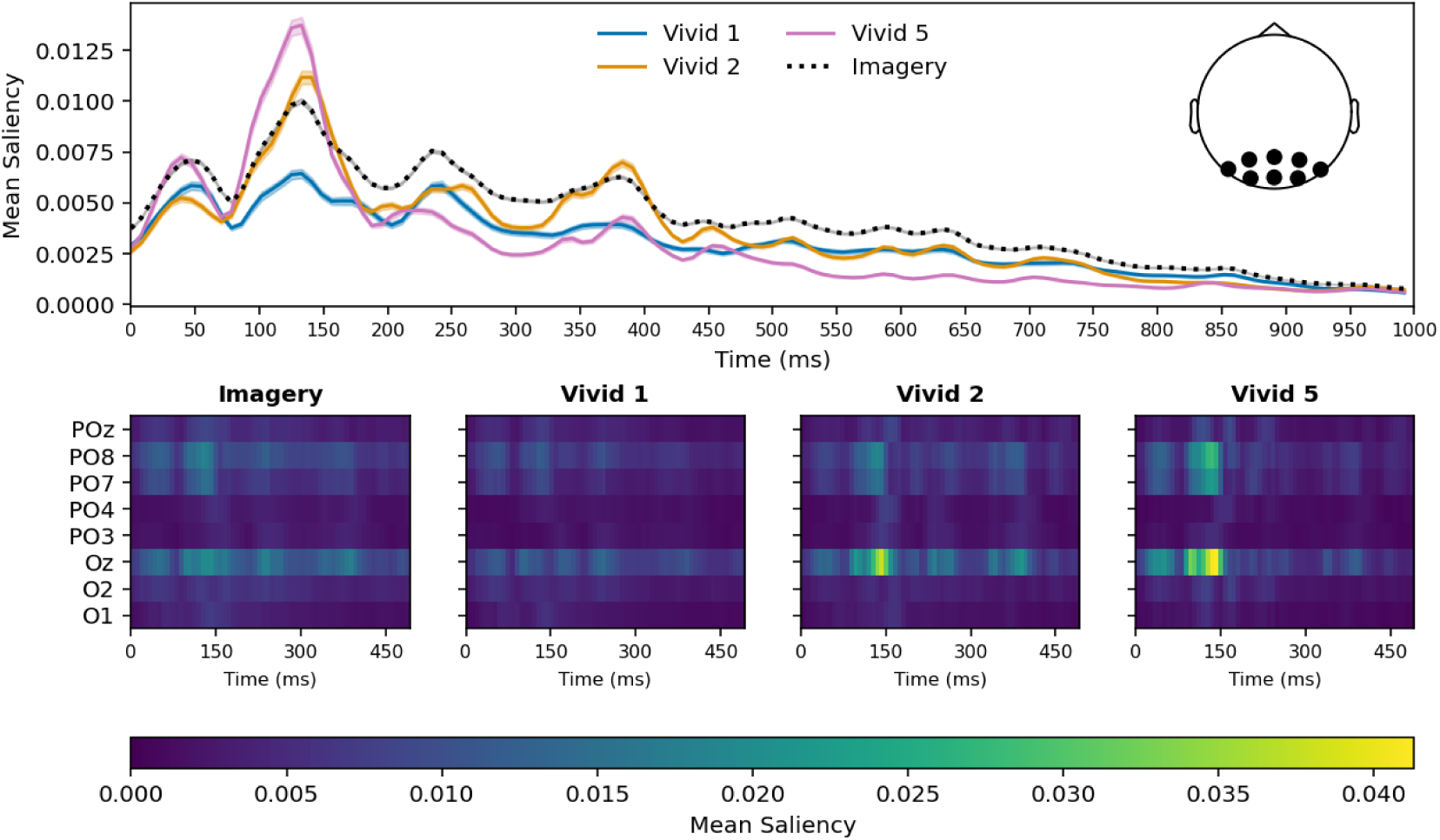
Mean temporal and spatial saliency maps for the 3-class EEGNet.

**Figure S6.**
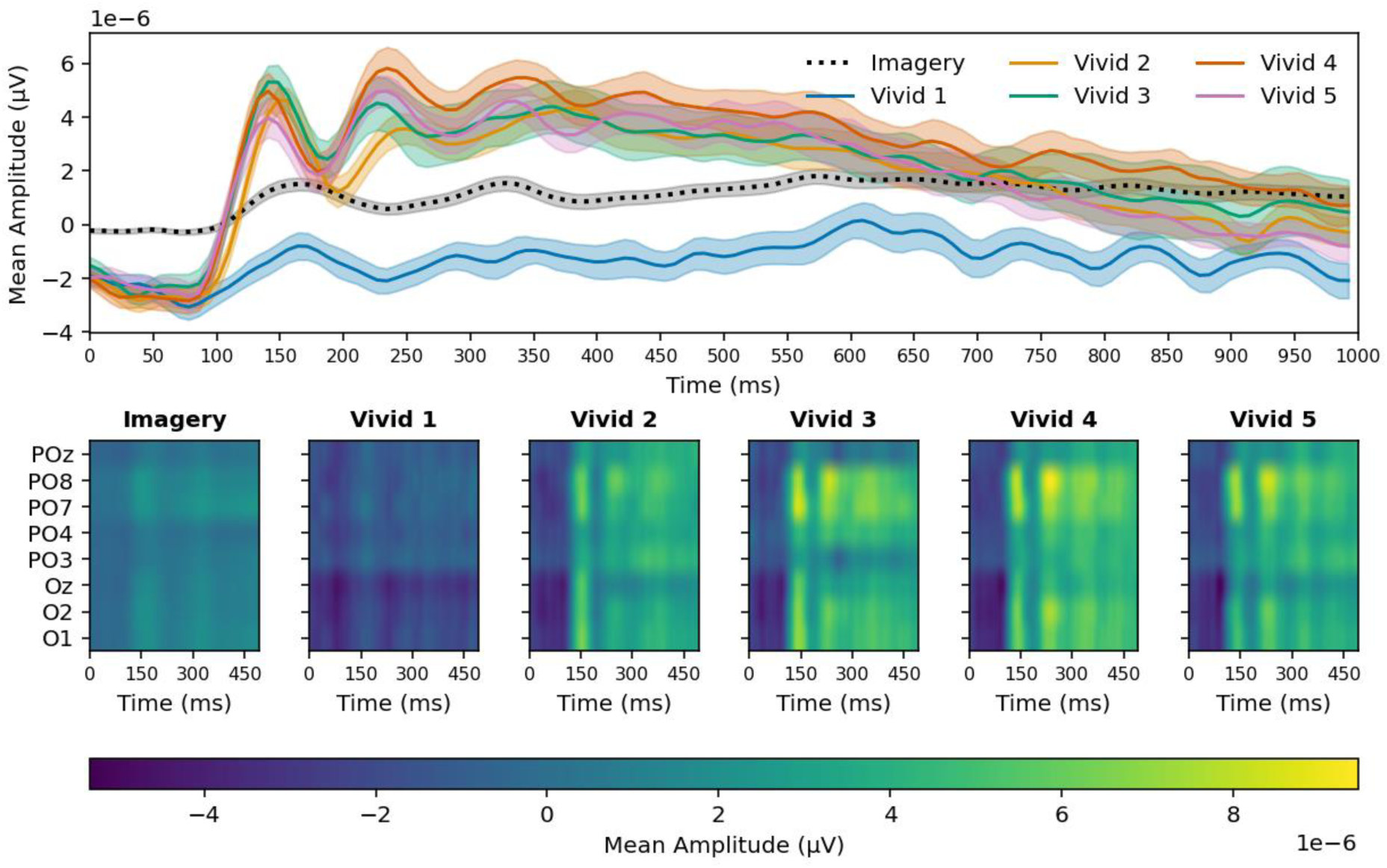
Grand-averaged Event-Related Potentials (ERPs) with SEM and mean amplitude per electrode across the first 500 ms. Bottom plot: Mean amplitude per electrode averaged across trials for the 0 to 500 ms time window. The fact that the patterns of activity are different from the saliency maps (Figure S4 & S5) suggests that EEGNet did not simply rely more on some electrodes because this is where the overall amplitude was the biggest, but because those electrodes contain the most relevant information for the classification task.

**Figure S7.**
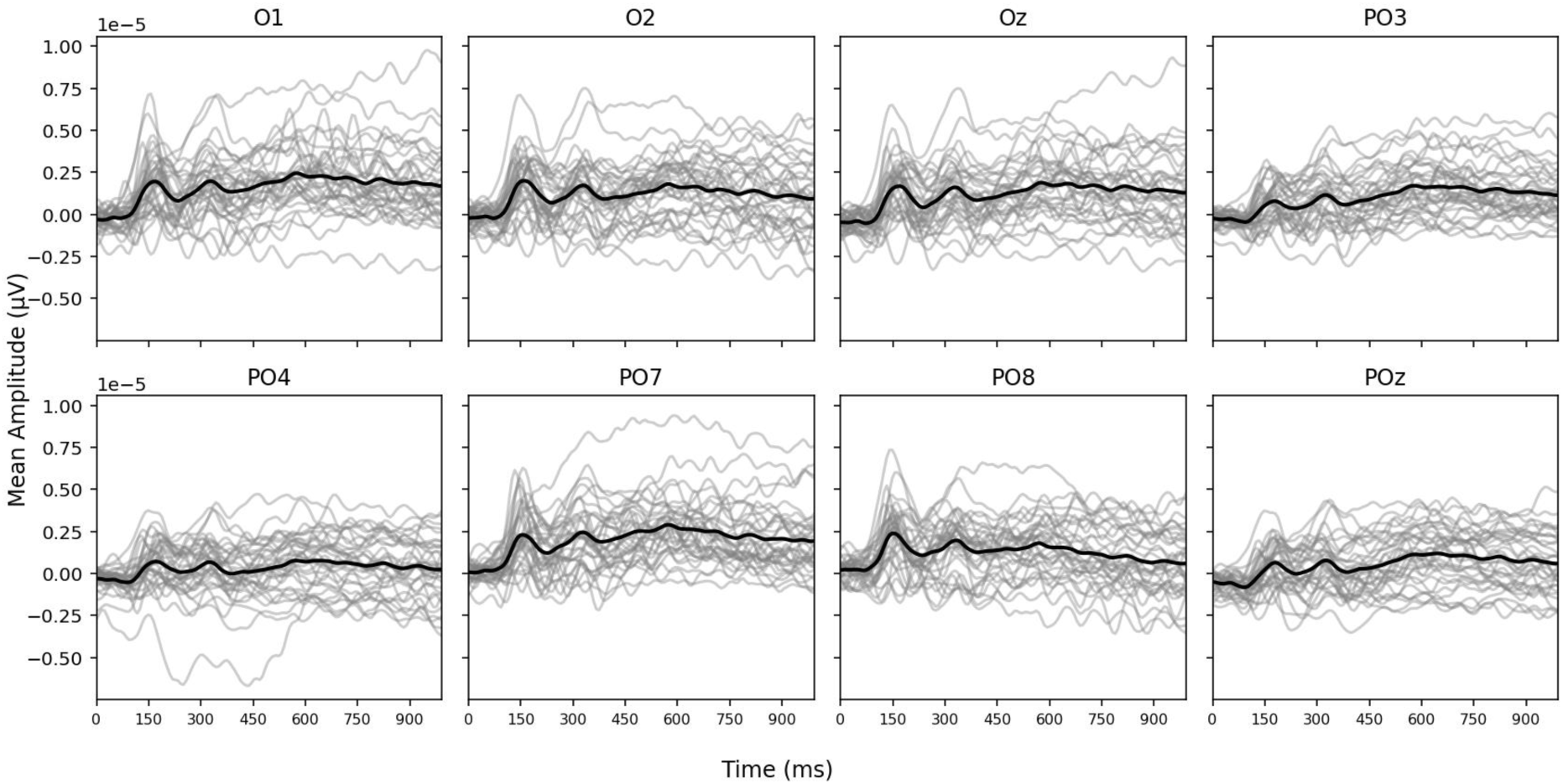
Grand-averaged and by-participant Event-Related Potentials (ERPs) for the imagery condition, for each of the 8 posterior electrodes. For each participant, all trials were averaged at each time point along the 0 to 1000 ms time window. The thick black line shows the grand-averaged ERP across the same time window.

**Figure S8.**
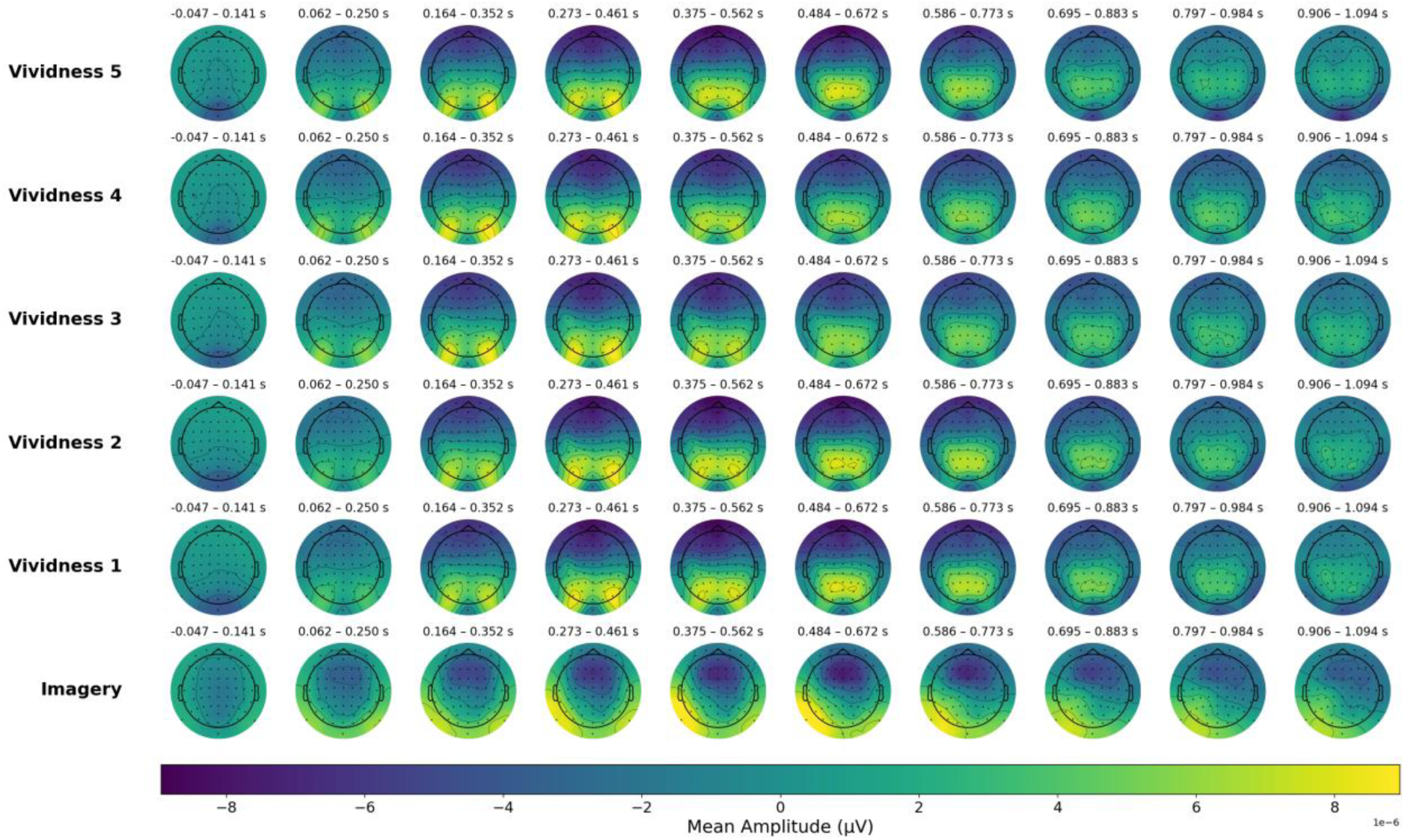
Time-resolved scalp topographies across vividness labels (perception) and imagery. Grand-average EEG scalp topographies are shown for each perceptual vividness level (class 1 to 5) and the imagery condition across ten time windows spanning -50–1050 ms post-stimulus. Each topography reflects mean activity averaged across trials and participants within a ∼200 ms moving window (step: ∼100 ms). Rows correspond to the vividness levels of the stimuli shown during the perception task (ordered from highest to lowest) and to the imagery task (bottom row).

## 3. Cats versus Strawberries control checks

The cat and strawberry reference images underwent the same image processing steps in order to compute their alternative, lower-vividness versions. Although both categories were regrouped under the same vividness labels for classification, physical differences in the original stimuli can persist in their subsequent derivative vividness levels.

**Does mean <u>accuracy</u> differ between cats and strawberries?**

- **Perception:**

- Participants: t(33) = -1.47, *p* = 0.150, *d* = 0.11, BF10 = 0.492
- 5-class EEGNet: t(33) = -3.57, *p* = 0.001, *d* = 0.60, BF10 = 28.7

**Does mean <u>vividness</u> differ between cats and strawberries?**

- **Perception**:

- Participants (reported): t(33) = -2.26, *p* = 0.030, *d* = 0.13, BF10 = 1.71
- 5-class EEGNet (expected): t(33) = -1.89, *p* = 0.067, *d* = 0.17, BF10 = 0.9

- **Imagery:**

- Participants (reported): t(33) = -1.95, *p* = 0.059, *d* = 0.10, BF10 = 0.995
- 5-class EEGNet (expected): t(33) = -1.55, *p* = 0.130, *d* = 0.15, BF10 = 0.545

### Interpretation

Despite potential physical differences between cats and strawberries that may have led to greater confusions for cats, EEGNet nonetheless successfully learned to classify vividness above chance for both categories (Cats: *M* = 42.41%, *SD* = 5.02, range 30% - 51.33%; Strawberries: *M* = 46.65%, *SD* = 8.64, range 32% - 63.33%; chance = 20%).

## Supplementary figures

**Figure S9.**
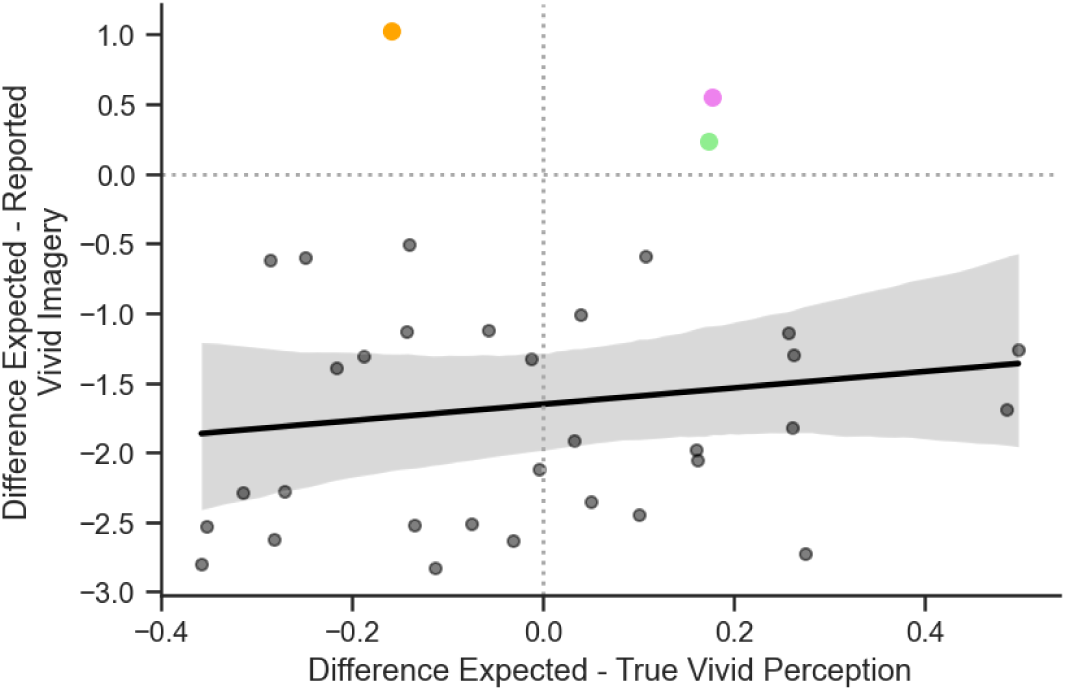
Mean by-participant difference between expected and reported (imagery) or labelled (perception) vividness, for the 5-class EEGNet.

**Figure S10.**
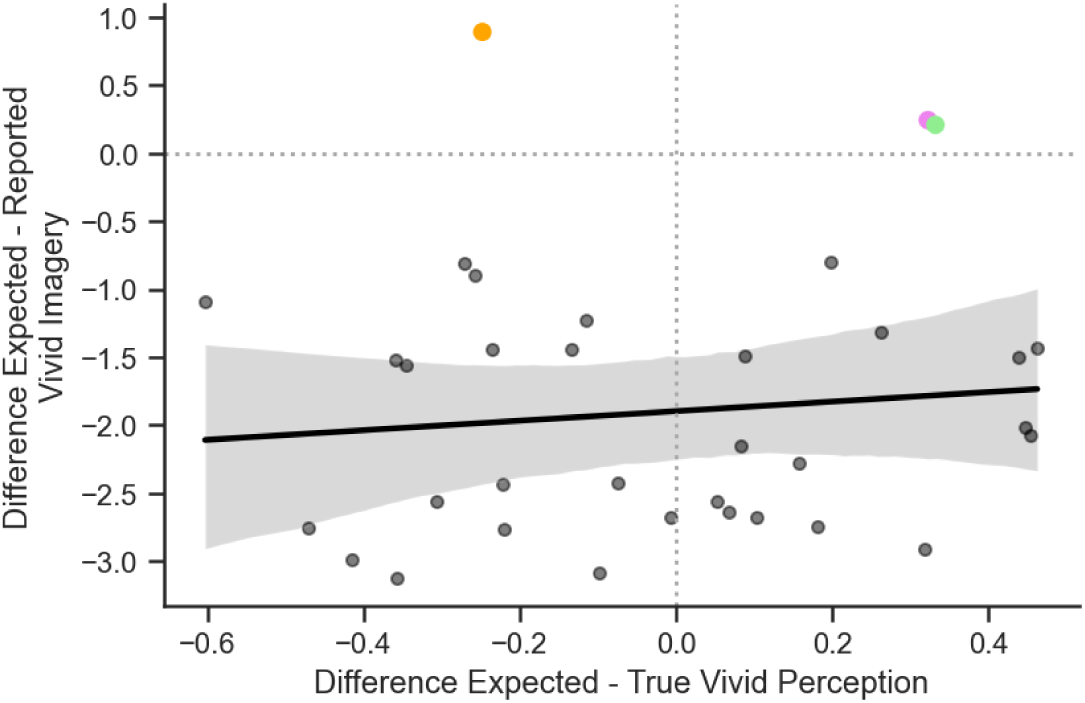
Mean by-participant difference between expected and reported (imagery) or labelled (perception) vividness, for the 3-class EEGNet.

## Supplementary Materials – Instruction screens

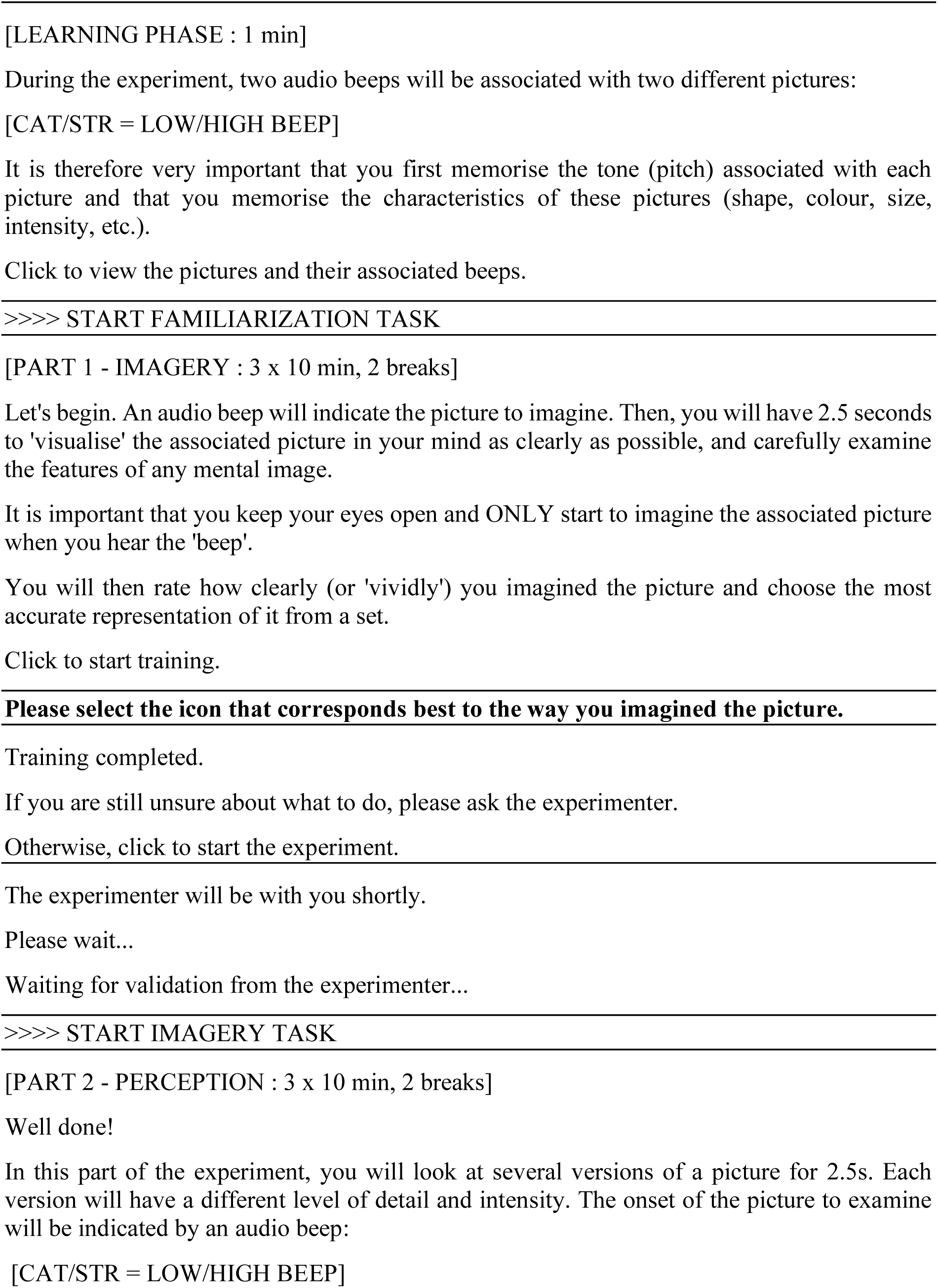
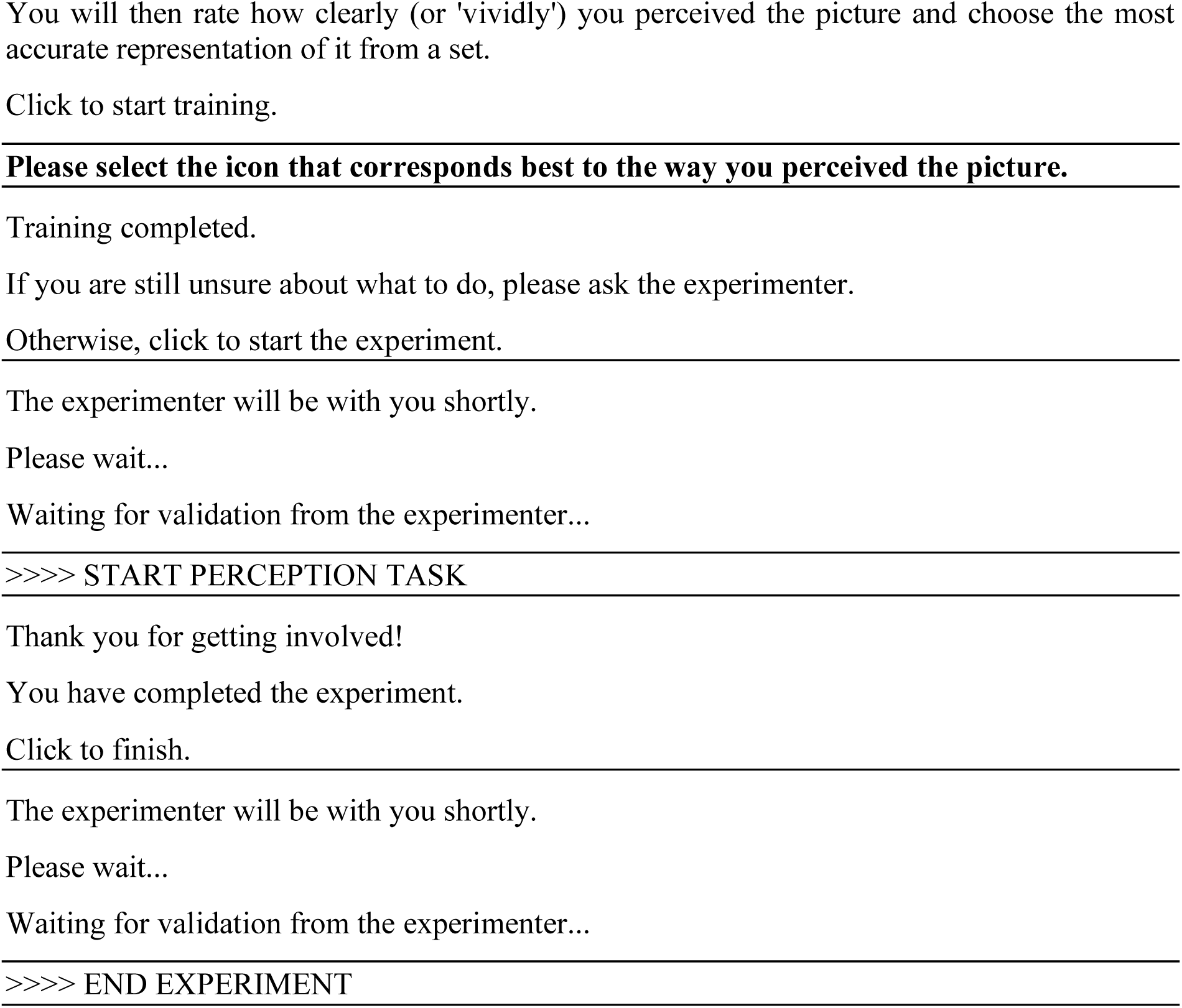

